# Large-pore connexin hemichannels function as molecule transporters independently of ion conduction

**DOI:** 10.1101/2024.02.20.581300

**Authors:** Pablo S. Gaete, Deepak Kumar, Cynthia I. Fernandez, Juan Manuel Valdez-Capuccino, Aashish Bhatt, Wenjuan Jiang, Yi-Chun Lin, Yu Liu, Andrew L. Harris, Yun L. Luo, Jorge E. Contreras

## Abstract

Connexin hemichannels were identified as the first members of the eukaryotic large-pore channel family that mediate permeation of both atomic ions and small molecules between the intracellular and extracellular environments. The conventional view is that their pore is a large passive conduit through which both ions and molecules diffuse in a similar manner. In stark contrast to this notion, we demonstrate that the permeation of ions and of molecules in connexin hemichannels can be uncoupled and differentially regulated. We find that human connexin mutations that produce pathologies and were previously thought to be loss-of-function mutations due to the lack of ionic currents are still capable of mediating the passive transport of molecules with kinetics close to those of wild-type channels. This molecular transport displays saturability in the micromolar range, selectivity, and competitive inhibition, properties that are tuned by specific interactions between the permeating molecules and the N-terminal domain that lies within the pore — a general feature of large-pore channels. We propose that connexin hemichannels and, likely, other large-pore channels, are hybrid channel/transporter-like proteins that might switch between these two modes to promote selective ion conduction or autocrine/paracrine molecular signaling in health and disease processes.

## INTRODUCTION

Connexin proteins assemble as hexamers to form hemichannels that are inserted in the plasma membrane. The docking of two hemichannels in apposed cells forms a gap junction channel (GJC) that allows direct cytoplasmic communication. Connexin hemichannels were the first identified plasma membrane channels with unique properties that mediate both the flux of atomic ions and the permeation of small molecules like ATP and glutamate. Opening of undocked hemichannels has been extensively associated with numerous pathological outcomes, which suggested that in physiological conditions these hemichannels remain mainly silent (i.e., in a closed state) to maintain the cellular electrochemical gradient. Nevertheless, a critical role for undocked hemichannels in physiological processes via the release of cytosolic molecules mediating autocrine and paracrine roles has emerged (1–5). How ion fluxes and molecular permeation coexist to mediate both physiological and pathological outcomes is still unclear.

Structural studies have revealed that connexin hemichannels have a wide pore cavity, with all six highly-flexible N-terminal (NT) domains lining the pore in the cytosolic entrance and forming the narrowest part of the conduction pathway (6–11). Although a fully physiological closed state has not been clearly captured yet, some of the resolved structures revealed a conformation with a pore entrance that was narrowed to a diameter of ∼5-8 Å, which could reflect a conformation in which only ions but not molecules can permeate (9–11). Functional assays showing the permeation of different small molecules suggested that the fully-open hemichannel conformation for permeation of molecules has an estimated limiting diameter of around ∼12-15 Å (5), which is likewise wide enough to allow ion fluxes. This has been partially supported by some structural studies (6–8); however, the mechanisms involved in molecular transport through the pore are still largely unknown. The present understanding is that molecular permeation in connexin hemichannels should be well-correlated with the movement of atomic ions, subject to size constraints, as suggested earlier for GJCs (12, 13). Under this rationale, the flux of both atomic ions and molecular permeants should proportionally increase, since a large pore size would act as a major determinant for both ion conductance and molecule permeation.

This notion, however, is inconsistent with findings showing uptake of fluorescent small molecules through connexin hemichannels (and the large-pore channels formed by pannexins) when there are negligible or undetected ionic currents, in particular at resting membrane potentials (14–19). Furthermore, pharmacological blockade of connexins or pannexins has been shown to differentially affect atomic ion and molecule permeation (15, 16). Given the importance of molecular signaling in physiology and pathology, the lack of understanding of this anomalous permeation phenomenon has been a confounder in understanding the biological and pathological roles of connexin hemichannels as well as for other large-pore channels.

To explore the mechanisms underlying the permeation of small molecules and ions in hemichannels, we evaluated the biophysical properties of permeation of two small fluorescent cationic dyes, ethidium and DAPI, whose permeation has been extensively used for assessing connexin hemichannel activity. By combining MD simulations, dye uptake assays, mutagenesis, cross-linking and electrophysiology, we found that permeation of molecules through connexin hemichannels exhibits properties that include saturability, competitive inhibition, selectivity, and discrete binding sites for molecules within the pore, reminiscent of classical transporter properties. We demonstrate that while ions and molecules use the same permeation pathway, the transport of molecules can be uncoupled from ion fluxes. We propose that connexin hemichannels may adopt different pore conformations that allow NT rearrangements to work either as ion channel or transporter for molecules. This finding has profound implications for our understanding for how connexin hemichannels, and perhaps large-pore channels in general, function in health and disease.

## RESULTS

### Transport of molecules through connexin hemichannels is selective and saturable

To properly understand how ions and molecules permeate through the connexin hemichannel pore, we recently developed a TEVC/dye uptake assay that measures kinetics of transport of fluorescent dyes and ionic current simultaneously in a single cell (19, 20). First, we compared the permeability of two membrane-impermeant fluorescent dyes, DAPI and ethidium, in two closely-related connexin isoforms, Cx26 and Cx30 (85.4% sequence identity). In line with previous findings (19), we found that Cx26 hemichannels were permeable to DAPI but not to ethidium (Fig. 1A-B), indicating a clear molecular selectivity. DAPI uptake through Cx26 hemichannels was concentration-dependent and saturated in the mid micromolar range (Fig. 1C-D). In contrast, Cx30 hemichannels were permeable to ethidium but not to DAPI (Fig. 1E-F). Ethidium permeation through Cx30 hemichannels saturated in the low micromolar range (Fig. 1G-H). The Km values for transport of these molecules were connexin isoform-specific: 90.1 ± 22.0 µM for DAPI in Cx26 and 6.6 ± 2.0 µM for ethidium in Cx30. This indicates that molecular transport through these two hemichannels is highly selective due to specific selective interactions between the permeants and the connexin channel pores.

**Figure 1.**
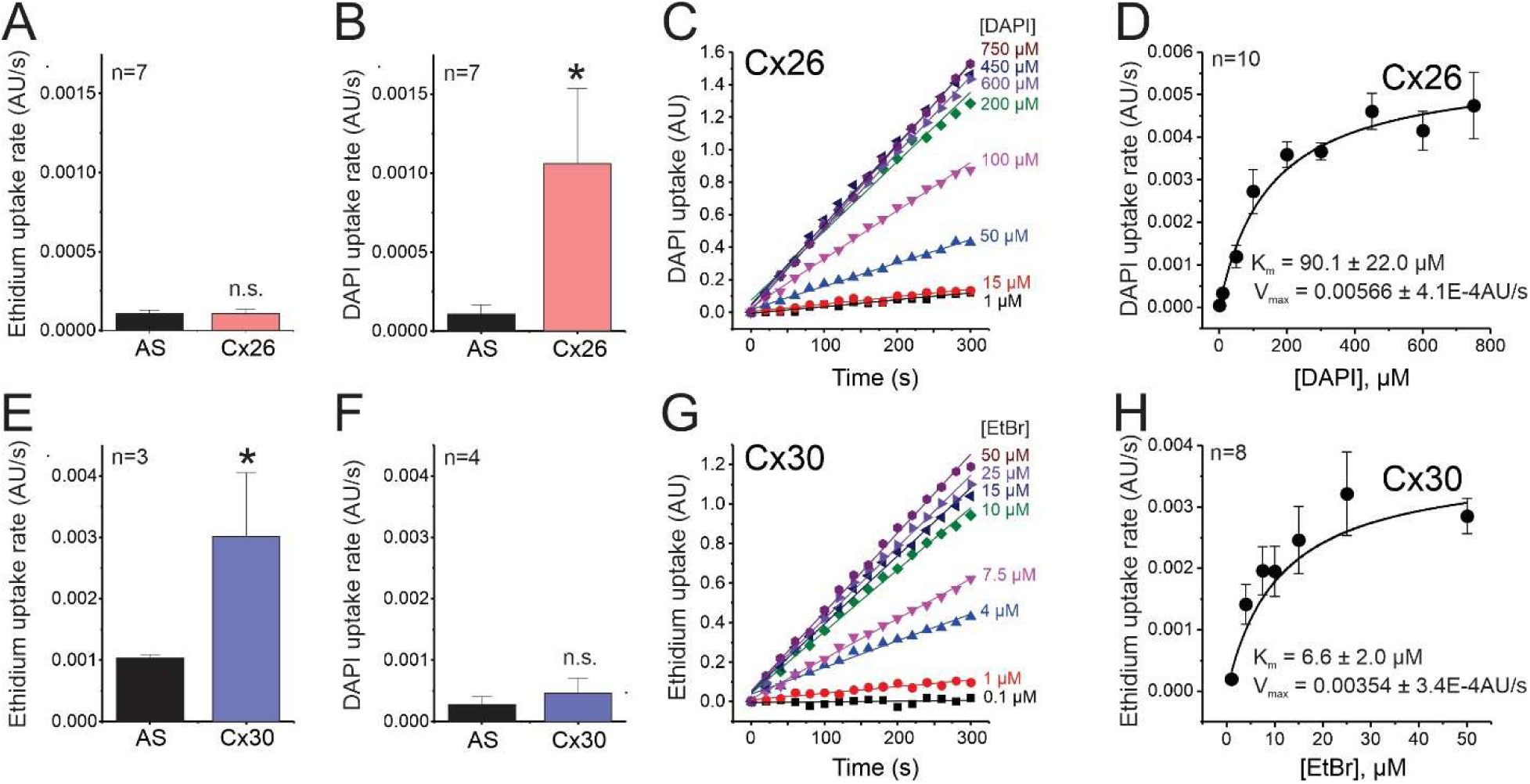
Transport of molecules through connexin hemichannels is selective and saturable. Dye uptake was evaluated in Xenopus oocytes expressing Cx26 (A-D) or Cx30 (E-H) at resting membrane potentials. Oocytes injected with antisense alone (AS) were used as controls (see Methods). **(A)** Ethidium uptake rate detected in oocytes incubated with 50 µM ethidium bromide (EtBr). **(B)** DAPI uptake rate detected in oocytes incubated with 50 µM DAPI dilactate. **(C)** Representative time courses of Cx26-mediated DAPI uptake. The concentration of DAPI dilactate in the extracellular bath is shown next to each trace. **(D)** Quantification of the DAPI uptake rate shown in **C**. **(E)** Ethidium uptake rate detected in oocytes incubated with 50 µM EtBr. **(F)** DAPI uptake rate detected in oocytes incubated with 50 µM DAPI dilactate. **(G)** Representative time courses of Cx30-mediated ethidium uptake. The concentration of ethidium bromide in the extracellular bath is shown next to each trace. **(H)** Quantification of the ethidium uptake rate shown in **G**. Data in **D** and H were fit with the Michaelis-Menten equation to calculate Km and Vmax. *, P < 0.05 vs AS by unpaired Student’s t-test. n.s.= non-significant. Error bars are SEM.

### Interactions between permeants and the NT domain determine selectivity and kinetics of molecule permeation

To gain insights into the mechanisms of molecule permeation through these hemichannels, we performed MD simulations. For this purpose, we used our equilibrated Cx26 hemichannel model, with which we have recapitulated experimental single-channel permeability of cAMP at ± 200 mV and Markovian milestoning simulations without voltage (21–23). Here, we examined qualitatively both ethidium (+1e) and DAPI (+2e) inward transport at -200 mV through the Cx26 hemichannel to identify potential protein-permeant interactions that may be responsible for the differences in permeability observed experimentally.

Three replicas of 500 ns all-atom simulations revealed two major ethidium binding regions in the Cx26 pore, one at the extracellular end and one at the NT region (roughly z = +20 Å and -10 Å, respectively) (Fig. 2A-B). The heightened probability density distribution in the extracellular region is attributed to electrostatic and van der Waals interactions with anionic side chain residues, including Glu42, Asp46, Glu47, and Asp50 (Fig. 2B and Supplementary Fig. 1). Within 500 ns, we observed complete permeation events in two of the replicates, suggesting that these interactions are short-lived (residence time 150-300 ns under -200 mV). In the NT region, however, ethidium exhibited a prolonged pi-stacking interaction with Trp3 sidechains in the third replica. This is consistent with our interaction analysis showing that the interaction between ethidium and Trp3 is the strongest among all pore-lining residues (Supplementary Fig. 1). We noticed that the relative orientation between ethidium and Trp3 sidechain plays a pivotal role in determining the access of ethidium through the Cx26 pore. The equatorial diameter of ethidium is 9.7 Å, smaller than the minimum pore diameter of 14 Å. Thus, ethidium can assume various orientations within the pore.

**Figure 2.**
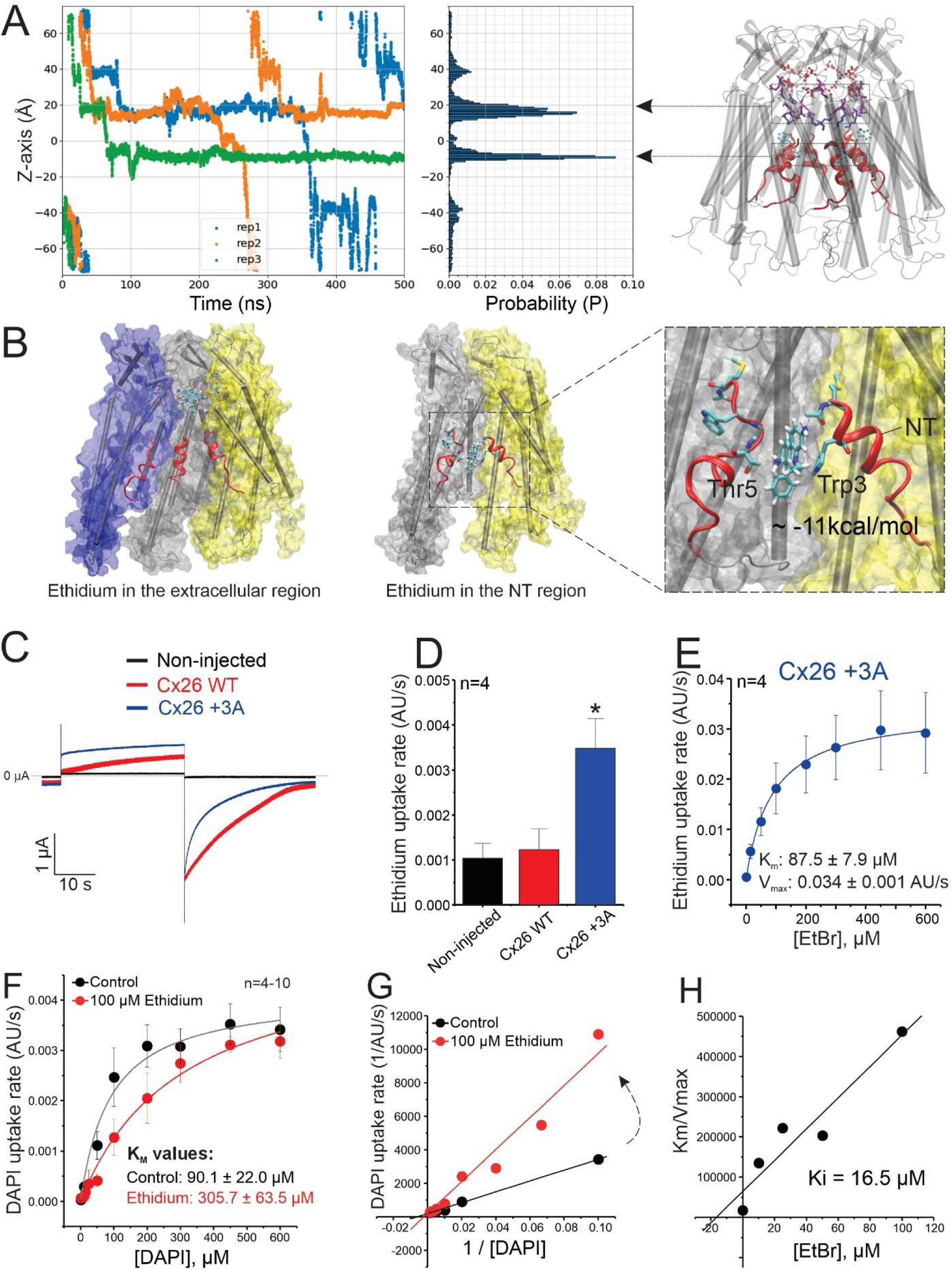
The N-terminal domain interacts with molecules and confers selectivity properties to connexin hemichannels. **(A)** MD simulations of ethidium permeation through Cx26 hemichannels. Traces represent the Z coordinates of the ethidium molecule. Flux is inward (from positive to negative Z coordinates) at −200 mV. Three independent replicas are shown in blue, orange, and green. The distribution probability analysis reveals two major interaction regions inside the pore. **(B)** Snapshot showing one ethidium molecule interacting in the extracellular region (left) and when trapped in the NT domain (middle). The total interaction energy for ethidium and Trp3 is shown (right). **(C)** Representative ionic current traces from oocytes expressing Cx26 wild-type (WT) or the mutant Cx26 +3A. Non-injected oocytes were used as controls. **(D)** Ethidium uptake rate recorded in oocytes incubated with 50 µM ethidium bromide (EtBr). **(E)** Concentration-response curve for ethidium uptake in oocytes expressing the mutant Cx26 +3A. **(F)** Effect of a saturating ethidium concentration on DAPI permeation. Experiments were performed in oocytes expressing Cx26 WT. **(G)** Lineweaver-Burk plot obtained from data shown in **F** reflects the competitive inhibition of ethidium on DAPI permeation. **(H)** Calculation of the inhibition constant (Ki) for ethidium. Data in **E** and **F** were fit with the Michaelis-Menten equation, to calculate Km and Vmax. Dye uptake was evaluated at resting membrane potentials. *, P < 0.05 vs non-injected, by one-way ANOVA plus Newman-Keuls post hoc test. Error bars are SEM.

When ethidium approaches the NT region of the pore horizontally, with its rings parallel to the membrane plane, the likelihood of establishing pi-stacking with Trp3 diminishes, while a vertical orientation of ethidium facilitates such interaction. When this interaction occurs, ethidium becomes trapped between Trp3 and the Thr5 from the neighboring subunit throughout the trajectory, presenting a basis for its impermeability (Fig. 2B).

Due to the limited sampling time and the stochastic nature of the non-equilibrium voltage simulations, the strong pi-stacking between ethidium and Trp3 was captured in only one replica. Hence, to experimentally verify if this is indeed the dominant interaction that impairs the permeability of ethidium in Cx26 hemichannels, we performed site-directed mutagenesis with the rationale of decreasing the ethidium affinity at this putative binding site. First, we inserted an alanine at position 3, next to the Trp3 (Cx26+3A). The mutant containing this insertion formed functional hemichannels with biophysical properties comparable to those of wild type, including similar gating properties, voltage-dependence, and sensitivity to extracellular Ca^2+^ concentrations, indicating that atomic ion permeation and global regulatory properties remain largely unchanged (Fig. 2C, and Supplementary Fig. 2). More importantly, the Cx26+3A mutation enabled ethidium permeation (Fig. 2D). A concentration-response analysis revealed that the transport of ethidium through this mutant hemichannel was saturable with a Km = 87.5 ± 7.9 µM (Fig. 2E). This Km is similar to that of DAPI in WT Cx26, suggesting the binding affinity of ethidium in Cx26+3A is in the same range as that of DAPI in WT Cx26. The data suggest that with the insertion of the Ala, Trp3 was either displaced deeper into the pore (further away from the Thr5 of the adjacent subunit) or reconfigured so that it could not form pi-pi interactions with ethidium. To distinguish these possibilities, Trp3 was substituted by alanine (W3A). This mutation also enabled ethidium permeation, and to the same degree as the Cx26+3A, showing that indeed the interaction of ethidium with Trp3 is what renders Cx26 impermeable to ethidium. Unexpectedly, the W3A mutation, in addition to enabling ethidium permeation, eliminated ionic currents (Supplementary Fig. 3). This striking initial finding is explored further in later experiments.

If ethidium has access to the WT Cx26 hemichannel pore and remains trapped at the NT-lined pore region, as predicted by MD simulations and supported by experiments, then its presence in the pore should decrease DAPI permeation in a concentration-dependent manner, since DAPI must traverse the same region to permeate. Consistent with this, we found that the apparent affinity for DAPI is reduced in the presence of 100 μM ethidium (Fig. 2F). Interestingly, a Lineweaver-Burk plot indicated that ethidium acts as a competitive inhibitor for DAPI permeation (Fig. 2G). A Km/Vmax relationship was obtained for DAPI permeation in the presence of different ethidium concentrations to estimate the inhibition constant (Ki). The Ki value for DAPI transport inhibition by ethidium was ∼16.5 µM (Fig. 2H). These data suggest that ethidium competes with or interferes with access to the DAPI binding site within the pore.

Voltage simulations were conducted to identify DAPI-pore interactions. DAPI (+2e) is a long and rigid molecule with +1e at each end. Its equatorial distance of 13.6 Å is comparable to the minimum pore diameter at the NT region. Although larger than ethidium, six permeation events were observed during three replicas of 500 ns DAPI simulations (Fig. 3A). In all replicas, DAPI established consistent interactions with Cx26 at several locations inside the pore (Fig. 3A and 3C). At the extracellular entrance, DAPI lays horizontally and interacts with the Asp50 sidechain of one subunit, and Asp46 and Glu47 from the opposite subunit (Fig. 3B, inset 1). This is followed by breaking of the interaction with Asp50 to form a new interaction with Glu42 in the same subunit (Fig. 3B, inset 2). Next, DAPI breaks the interaction with Glu47 and Glu42 and forms new interactions with two Asp2 in the neighboring NT helices (Fig. 3B, insets 3 and 4). However, the DAPI-Asp2 interaction has shorter lifetime (< 100 ns) than ethidium-Trp3 interaction (> 400 ns). Breaking the charge-charge interaction with one of the Asp2 residues reorients DAPI vertically in the pore, which allows DAPI to pass through the narrowest pore region (Supplementary Fig. 4). To confirm DAPI’s interaction with Asp2 at the end of the NT domain, we replaced residue Asp2 with asparagine (D2N). As expected, the D2N mutation was still permeable to DAPI, but it lowered the apparent affinity ∼ 3-fold without affecting the maximum transport velocity (Fig. 3D). Altogether, our *in silico* and functional data demonstrate that DAPI and ethidium interact with the NT domain through different types of interactions, and selectivity and transport kinetics are determined by these interactions.

**Figure 3.**
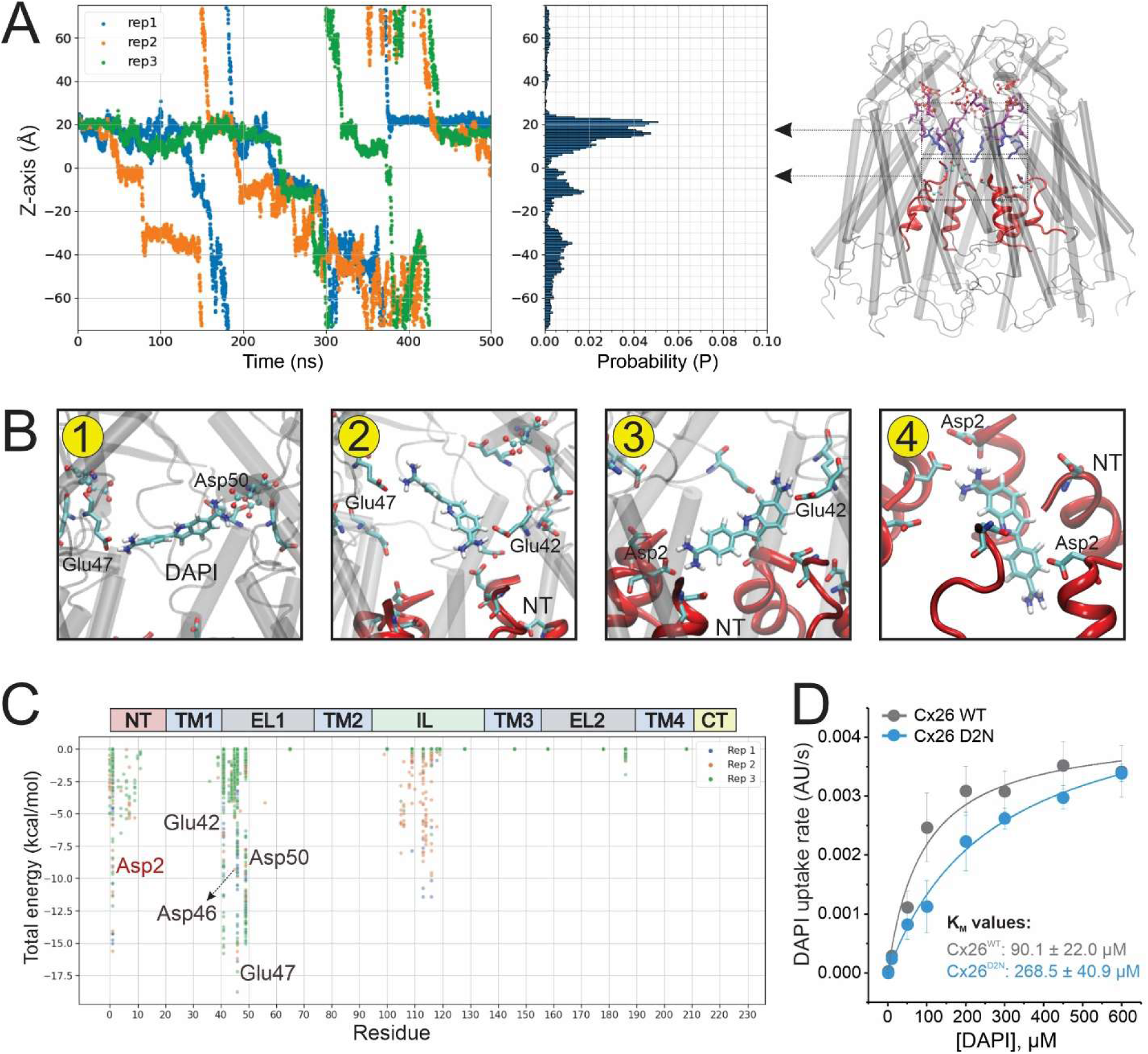
Interactions between permeants and the N-terminal domain determine the kinetics of molecular permeation. **(A)** Simulations of DAPI permeation through Cx26 hemichannels. Traces represent the Z coordinates of the DAPI molecule when the flux is inward (under −200 mV). Three independent replicas are shown in blue, orange, and green. A snapshot of the simulated system is shown in the right panel. The protein backbone is shown using gray cylindrical helices in cartoon mode. N-terminal (NT) domains are highlighted in red. **(B)** One DAPI molecule and the sidechains interacting with DAPI during a single permeation event are shown in licorice with atom color code (red, oxygen; cyan, carbon; blue, nitrogen; white, hydrogen). Lipids, ions, and water molecules are not shown for clarity. Panels 1-4 represent the interactions between DAPI and the residues lining the pore during the permeation process. Inset 1: DAPI interaction in the extracellular region of the pore; inset 2: DAPI translocation to the NT region; inset 3: DAPI interaction with the NT domain; inset 4: DAPI translocation to the intracellular region (see Results for detailed description). Transmembrane domains are not shown in inset 4 for clarity. **(C)** An MMGB pairwise analysis was performed to calculate the total interaction energies between DAPI and the resides lining the pore of Cx26 hemichannels. **(D)** The effect of D2N mutation on DAPI permeation unveils the importance of the NT in the control of molecule permeation kinetics. DAPI uptake was evaluated at resting membrane potentials. Error bars are SEM.

### NT domain rearrangements are necessary for permeation of molecules

The narrowest pore region in Cx26 hemichannels is located at the end of the NT domain, where these molecules are interacting. Therefore, we asked whether the dynamics of NT domain are coupled with molecular permeation. In all three replicas of 500 ns simulations, DAPI interaction with a single NT domain led to a slight unfolding of the interacting NT and pulled it toward the cytoplasmic entrance, reflected in the distance between Met1 of adjacent subunits (Supplementary Fig. 5).

This rearrangement was coupled with a loss of interactions between the NT domain and the TM1 helix, including a salt bridge between Asp2 and Lys41, and hydrophobic interactions between Trp3-Met34 and Trp3-Ile33 (Supplementary Fig. 6). These observations suggest that permeation of DAPI through the pore exerts effects on the conformational dynamics of the NT. Hence, to assess whether conformational rearrangements of the NT are necessary for permeation of molecules, we created a mutant channel with a cysteine inserted in the NT domain at position 2 (named Cx26B +2C, Fig. 4A, and Supplementary Fig. 7). In this mutant, residues C211 and C218 were replaced with serine to prevent unspecific effects. Thus, hemichannels including C211S and C218S but not the cysteine at position 2 (named Cx26B) were used as control. We then assessed the rate of disulfide bridges formation between adjacent cysteines in state-dependent manner in the presence of an oxidant agent, TBHO_2_. For this, we used the well-known ability of millimolar extracellular Ca^2+^ to induce a closed state of the hemichannels (24). First, we showed that the Cx26B +2C mutant hemichannels displayed similar Ca^2+^-gating sensitivity when compared to wild-type hemichannels (Supplementary Fig. 7). Figure 4B shows representative ionic current traces for Cx26B and Cx26B +2C in the presence of 0.1 mM and 5 mM Ca^2+^, two conditions that favor opening and closing of hemichannels, respectively. When 2 mM TbHO_2_ was perfused to promote formation of disulfide bridges, a mono-exponential decay of the ionic currents was observed in the Cx26B +2C, but not in the Cx26B hemichannels, suggesting that formation of a disulfide bond at position 2 blocked ionic currents. Notably, the current decay induced by TbHO_2_ was accelerated as Ca^2+^ concentrations were increased suggesting the likelihood of disulfide linking between cysteines occurs more readily when channels are in a physiological closed conformation (Fig. 4B-C). Disulfide bridge formation was further confirmed via connexin dimer generation in non-reducing western blot analysis (Supplementary Fig. 8). In addition, we evaluated whether formation of metal bridges with Cd^2+^ (100 nM) mimicked the effect observed with TbHO_2_. As expected, the exponential current decay by Cd^2+^ blockade was also accelerated in a Ca^2+^-dependent manner (Fig. 4D). Taken together, these data suggest that the end of the NTs (where +2C was introduced) come in close proximity when hemichannel transitions with higher occupancy to the closed state induced by extracellular Ca^2+^.

**Figure 4.**
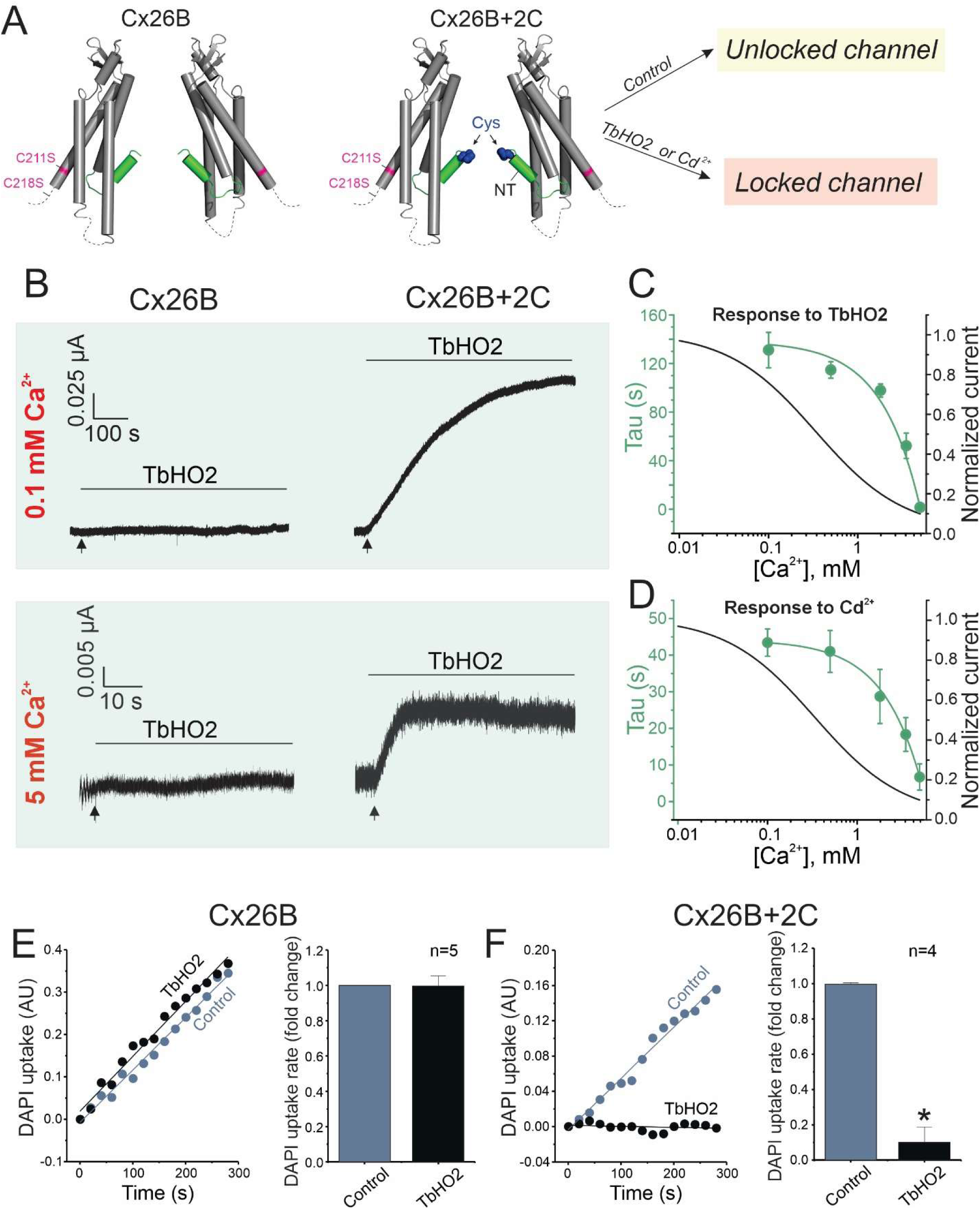
Molecules require NT rearrangements for translocation through the Cx26 hemichannel. **(A)** Scheme of mutants used for the crosslinking approach. For clarity, only two connexin monomers are shown. In the mutant Cx26B +2C, one cysteine (in blue) was inserted at position 2 in the NT domain, shown in green. Additionally, residues C211 and C218 (in magenta) were replaced with serine to prevent cross-linking at those locations. Hemichannels formed by C211S and C218S without the cysteine at position 2 (named Cx26B) were used as controls. In theory, only Cx26B +2C channel pore can be locked by the formation of disulfide bridges, when the cysteines are exposed to oxidant agents, such as TbHO_2_ or by formation of metal bridges, when cysteines are exposed to Cd^2+^. **(B)** Representative traces of ionic current recorded in oocytes expressing Cx26B or Cx26B +2C. Oocytes were perfused with Ringer solution containing specific Ca^2+^ concentrations, and membrane potential was held at -40 mV. After ionic current stabilization, oocytes were perfused with 2 mM TbHO_2_ to lock the channels by crosslinking. Arrows indicate the time TbHO_2_ perfusion starts. **(C)** Analysis of the kinetics of disulfide-bridge formation by calculation of tau values after TbHO_2_ perfusion. **(D)** Analysis of the kinetics of metal-bridge formation by calculation of Tau values after perfusion with 100 nM Cd^2+^, following the same experimental approach shown in **B-C**. Note that channel closure by TbHO_2_ or Cd^2+^ is faster when the relative open probability is low, and vice versa. **(E)** Time course of DAPI uptake in oocytes expressing Cx26B (control) before and after incubation with 2 mM TbHO_2_. For quantification, the DAPI uptake rate was normalized to the control group. **(F)** Time course and quantification of DAPI uptake in oocytes expressing Cx26B +2C before and after incubation with 2 mM TbHO_2_. For **E** and **F**, DAPI uptake was evaluated in oocytes held at -40 mV. *, P < 0.05 vs control by paired Student’s t-test. Error bars are SEM.

Next, we applied the same strategy to determine whether molecule permeation is affected by the crosslinking between cysteines in the NT. In control conditions, hemichannels formed by Cx26B +2C were permeable to DAPI as observed in wild type hemichannels. However, upon treatment with 2 mM TBHO_2_ DAPI permeation was completely abolished in Cx26B +2C hemichannels, but not in the control Cx26B hemichannels (Fig. 4E-F). This finding indicates that atomic ions and molecules are transported through the pore and that NT rearrangements are necessary for both ionic and molecule permeation.

### Transport of molecules occurs independently of ionic conduction

Previous studies reported high permeability rates for fluorescent dyes at negative membrane potentials, while ionic currents remain undetected (14, 18). These findings contradict the concept that a channel with an intrinsically wide permeation pathway will permit free diffusion for both ions and molecules. Taking advantage of the TEVC/dye uptake assay, we assessed whether molecule permeation correlates with the increase of ionic flux in a voltage-dependent manner. DAPI uptake in oocytes expressing Cx26 was detected when membrane potential was held at negative membrane potentials (Fig. 5A). Strikingly, DAPI uptake was abolished at positive membrane potentials at which the ionic conductance was robust (Fig. 5A). The voltage dependence of DAPI transport displayed a Boltzmann relationship inverse to the relative open probability for ionic currents (Fig. 5B). This finding strongly indicates that DAPI and atomic ions are not moving simultaneously within the pore.

**Figure 5.**
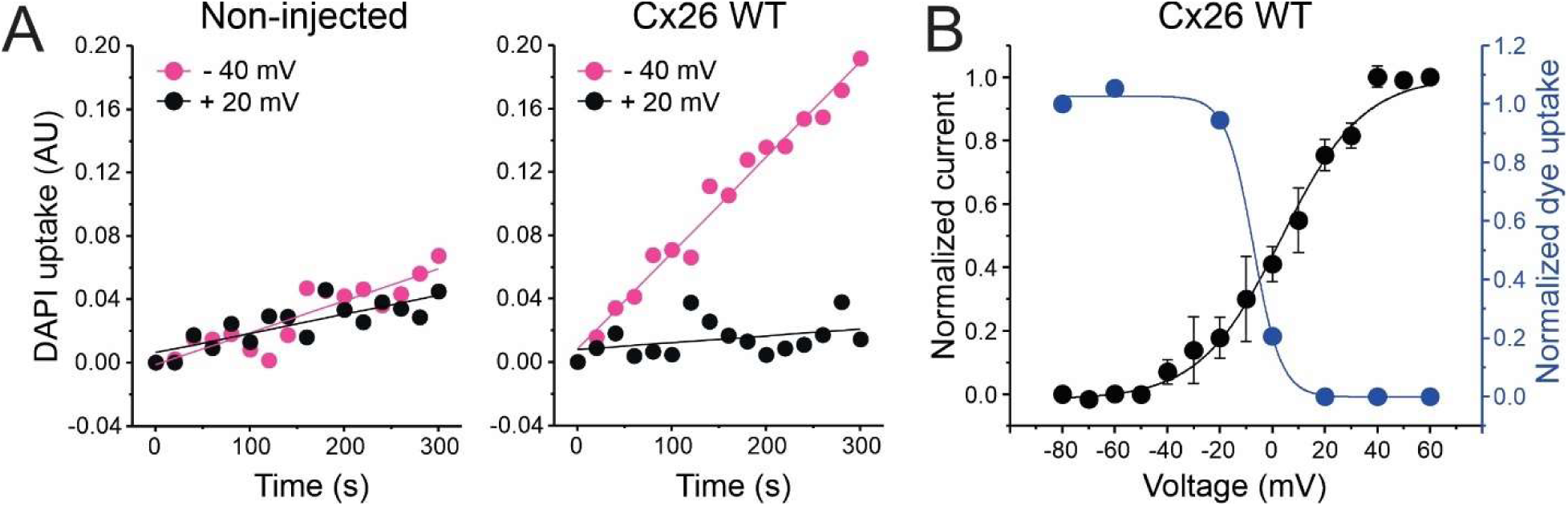
The transport of molecules in connexin hemichannels is voltage-dependent, and favored at physiological negative membrane potentials. **(A)** Time course of the DAPI uptake evaluated in non-injected oocytes (left) or oocytes expressing Cx26 WT (right). Oocytes were perfused with Ringer solution containing 1 mM Ca^2+^ plus 50 µM DAPI dilactate and membrane potential was clamped at -40 mV (magenta). After 5 min recording, membrane voltage was switched to +20 mV (black) and DAPI uptake was recorded again. **(B)** Effect of membrane potential on ionic current (black) and DAPI permeation rate (blue) through Cx26 hemichannels. Curves were fit with the Boltzmann equation. Error bars are SEM.

If the flux of molecules and ions occurs simultaneously within the same conduction pathway, it would be expected that molecules within the pore should reduce the atomic ion flux. Thus, we tested whether macroscopic currents detected by TEVC in oocytes expressing Cx26 are affected by the presence of a saturating concentration of DAPI (200 µM) in the extracellular bath. We observed that DAPI did not affect the macroscopic ionic currents in Cx26 hemichannels (Fig. 6A). Consistently, single-channel conductance was not affected by the presence of DAPI in the pipette. This suggests that molecules while using the same pore than atomic ions, do not use it simultaneously suggesting a different mechanism of permeation. (Fig. 6B-C).

**Figure 6.**
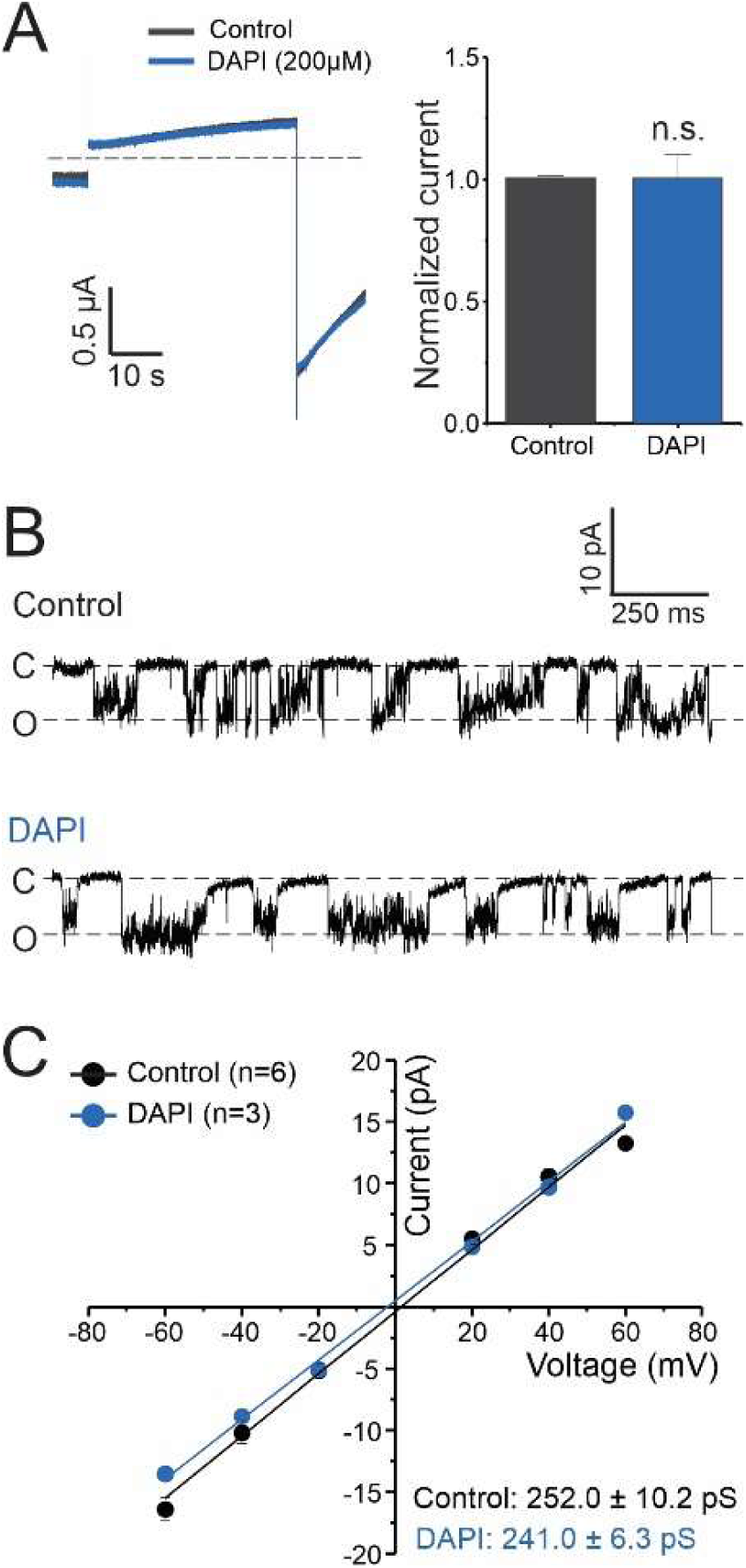
Permeant molecules do not affect macroscopic ionic currents or single-channel conductance of Cx26 hemichannels. **(A)** Representative ionic current traces recorded by TEVC in oocytes expressing Cx26 WT. Opening of Cx26 hemichannels was induced by a 40 s depolarizing pulse (see Methods). A saturating concentration (200 μM) of DAPI was added to the extracellular bath to determine the effect of this permeant on atomic ion permeation. The quantification of the magnitude of tail currents is shown in the bar graph. Values were normalized to the control group. **(B)** Representative single-channel traces recorded in the cell-attached configuration in oocytes expressing Cx26. Dashed lines show the closed (C) and open state (O). **(C)** Quantification of current-voltage relationship from single channel recordings. Single channel conductance for each group is shown. n.s.= non-significant vs control by paired Student’s t-test. Error bars are SEM.

To unequivocally confirm that molecules can translocate independently of atomic ion flux and building on the striking previously mentioned findings for the Cx26^W3A^ mutant (Supplementary Fig. 3), we revisited two human NT mutations that previously were shown to lack ionic currents (i.e., loss-of-function mutations): Cx26^N14Y^ and Cx26^S17F^ (25, 26). First, we confirmed by TEVC that these mutations render Cx26 hemichannels ‘silent’ for ion fluxes (Fig. 7A-C and Supplementary Fig. 9A-B), and that their expression levels at the plasma membrane were unaffected by the mutations (Supplementary Fig. 9C-D). Then, we tested if these ‘silent’ hemichannels could transport molecules. Notably, we observed both mutant channels retained the capability to transport DAPI when expressed in Xenopus oocytes or HeLa cells (Fig. 7D-F, and Supplementary Fig. 10). Furthermore, these channels were also permeable to ethidium (Supplementary Fig. 11). Thus, these mutations at the NT block ionic conductance, but permit the WT permeability to DAPI, and impart permeability to ethidium. Altogether, these data suggest that human mutations that blunt ionic currents do not necessarily affect transport of molecules, supporting that these processes are uncoupled.

**Figure 7.**
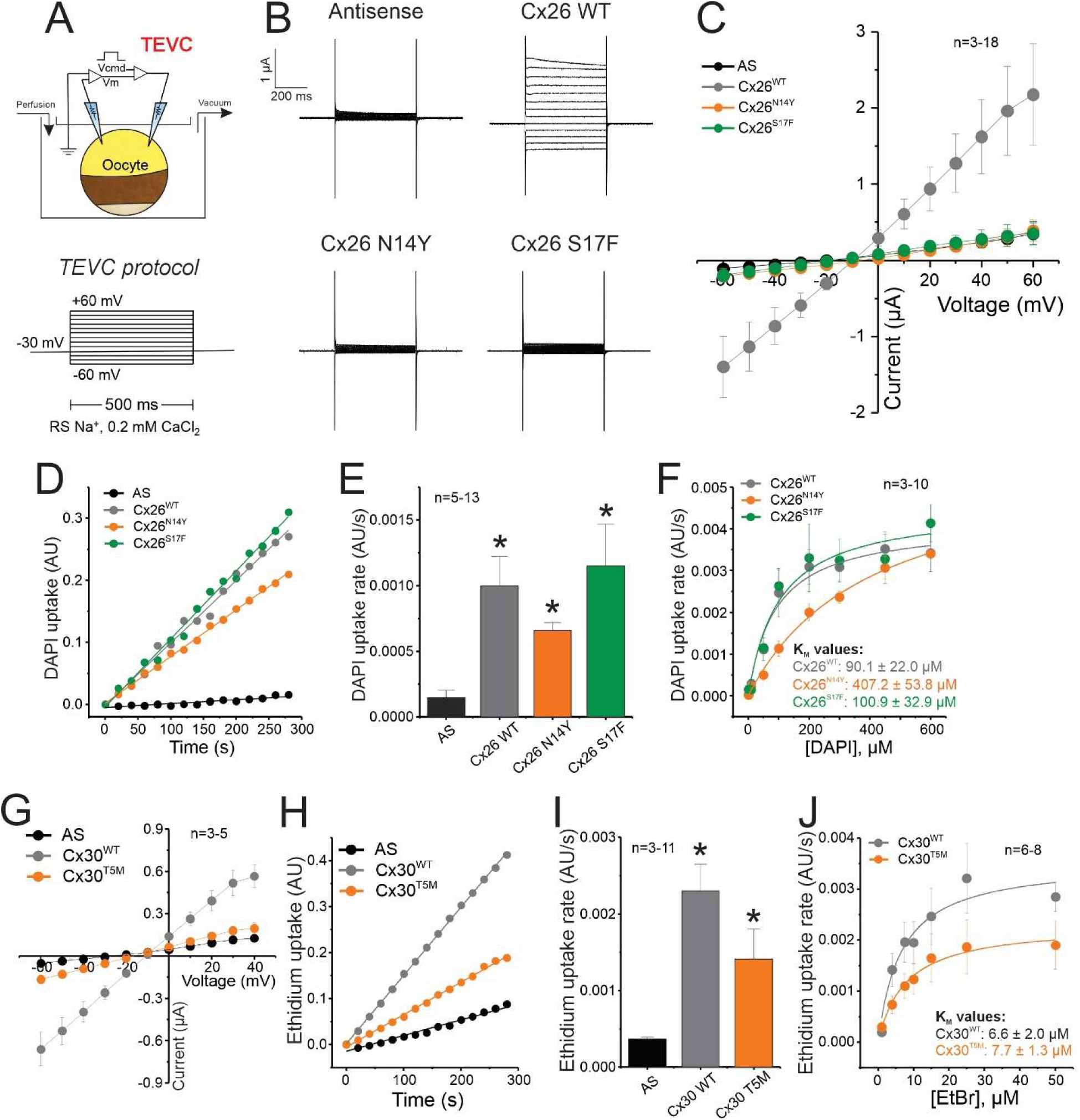
Pathologic human mutations in connexin hemichannels that lose atomic ion permeability retain the ability to transport small molecules. **(A)** Preparation scheme and TEVC protocol for experiments shown in **B**, **C**, and **G**. TEVC experiments were performed at low Ca^2+^ concentration to increase the relative open probability. **(B)** Representative traces of the ionic current observed in control oocytes injected with an antisense oligonucleotide (AS) or in oocytes expressing Cx26 mutants. **(C)** I-V relationship from data shown in **B**. **(D)** Representative time courses of DAPI uptake in oocytes expressing Cx26 mutants incubated with 50 µM DAPI dilactate. **(E)** Quantification of DAPI uptake rates from data shown in **D**. **(F)** Effect of N14Y and S17F mutations on DAPI permeation kinetics. **(G)** I-V relationship obtained from TEVC experiments performed in oocytes expressing Cx30 or the mutant Cx30^T5M^, using the same approach shown in panel **A**. **(H)** Representative time courses of ethidium uptake in oocytes expressing Cx30^T5M^ incubated with 50 µM ethidium bromide (EtBr). **(I)** Quantification of data shown in **H**. **(J)** Effect of T5M mutation on ethidium permeation kinetics. Dye uptake was evaluated at resting membrane potentials. *, P < 0.05 vs AS, by one-way ANOVA plus Newman-Keuls post hoc test. Error bars are SEM.

To test whether this observation is restricted to Cx26 hemichannels, we repeated the same approach with Cx30. The T5M mutation abolished the ionic currents (Fig. 7G). As observed with Cx26 mutants, hemichannels formed by Cx30^T5M^ were permeable to ethidium like the wild-type hemichannel (Fig. 7H-I). We observed a slight reduction in the distribution of Cx30^T5M^ at the plasma membrane and a decrease in the total protein (Supplementary Fig. 12), which correlated with a lower Vmax for ethidium flux (Fig. 7J), suggesting that the magnitude of the transport was reduced in oocytes expressing Cx30^T5M^, likely due to the reduction in the number of hemichannels at the plasma membrane. Notably, the Km value for ethidium transport in Cx30^T5M^ was identical to that from wild-type Cx30 hemichannels, indicating that the kinetic properties for molecular transport were unaffected by the T5M mutation (Fig. 7J). These data demonstrate that molecules are transported independently of atomic ion fluxes in at least two connexin isoforms.

## DISCUSSION

A general assumption about the permeation of molecules through large-pore channels is that molecule flux is by passive diffusion through the pore, along with the diffusing atomic ions, and that small molecules are discriminated mainly based on the limiting pore width and intrapore charge density when the channel is in the open state. Here, we explored the biophysical properties of the transport of cationic fluorescent dyes through connexin hemichannels to gain fundamental insights regarding the mechanisms that underlay molecule permeation in large-pore channels. While previous studies have shown that different connexin channels can have different molecular selectivity not readily explainable on the basis of size or charge (19, 27, 28), our findings point to the role of specific residues, primarily in the NT domain, that interact with permeants to impart a high degree of selectivity among molecules of similar size and charge. Further, we show that molecular transport throughout these channels is saturable in the micromolar range and displays competitive inhibition. These features are controlled by identified interactions within the pore, particularly with the NT domain. Most important, our findings are inconsistent with the current view that permeation of ions and molecules through large-pore channels occurs by the same mechanism. We provide evidence that permeation of molecules and of atomic ions can be uncoupled from each other and likely rely on different conformational states and/or dynamics.

### Permeation of molecules through connexin hemichannels: Saturability, Binding and Competition

Our data demonstrate that transport of molecules through Cx26 and Cx30 is saturable, and at surprisingly low concentrations, which is consistent with a facilitated diffusion mechanism. This property may be shared by other channels forming a wide pore; saturation of cationic fluorescent dye transport has also been found for Cx43 hemichannels and CALHM-1 channels (19, 29, 30). Saturability is not specific to cationic molecules since the flux of cAMP, a negatively charged endogenous permeant, is also saturable (Supplementary Fig. 13). Importantly, saturation for all tested molecules occurs in the micromolar range, indicating that saturation of molecule transport does not occur due to the crowding of molecules within the pore. Notably, the Km values calculated for molecule transport in connexin hemichannels are considerably lower than the Km reported for some GLUT transporters, where a millimolar concentration is needed to saturate the facilitated diffusion process (31, 32).

It has been suggested that the molecular selectivity of GJCs and hemichannels arises from binding sites within the pore (27, 33). Recent computational studies of cAMP permeation through Cx26 hemichannels suggested that the pore contains discrete binding sites that interact with cAMP (22). These findings predict saturability of molecular permeation, as we have now demonstrated. Indeed, the occupancy probabilities for DAPI and ethidium along the pore (Fig. 2A and Fig. 3A) suggest that both fluorescent dyes establish favorable interactions with residues at both the NT region and at the extracellular entrance of the pore (where the Ca^2+^ binding region is located). This is consistent with studies showing that these two regions are critical for molecular selectivity (16, 19).

The strongest interaction observed in the MD simulations was a pi-pi stacking between ethidium and residue Trp3 at the NT of Cx26, which hampered *in silico* permeability to ethidium. We experimentally demonstrated that the ethidium impermeability in Cx26 hemichannels likely resulted from this interaction because: (1) W3A mutation or insertion of an alanine at position 3 (+3A) to disrupt this pi-pi stacking interaction enabled ethidium transport; (2) ethidium acts as a competitive inhibitor for DAPI permeation, which indicates that ethidium has access to the inner pore, as predicted by MD simulations even though not permeable. This is also consistent with the finding from MD simulations and mutagenesis experiments showing that DAPI interacts with the NT domain, specifically with Asp2, in practically the same region as ethidium.

Importantly, the NT domain in connexin channels defines the narrowest and most flexible part of the pore as previously shown by cryo-EM, MD simulations and functional studies (6–11, 19, 22, 34, 35). In our MD simulations, including the one reported previously for cAMP, permeants rotate in response to the local charge and geometry of the pore as they are moving through (22). In particular, the NT serves as a common domain for molecule recognition through a combination of electrostatic, hydrophobic interactions, as well as the entropic effects associated with the different orientations and conformations of the molecule. Hence it is not surprising that mutations in the NT region often have significant impact on molecule selectivity and permeation kinetics, as demonstrated by the shift in the apparent affinity when we mutated these sites. For example, ethidium uptake was enabled with different mutations in the NT of Cx26, in particularly Cx26+3A (Fig. 2A), W3A (Supplementary Fig. 3), N14Y, S17F (Supplementary Fig. 11) and N14K (19).

### Uncoupling of molecule transport and atomic ion flux

More than 20 years ago, Contreras et al., showed that permeation of molecules can occur at resting membrane potentials in Cx43 hemichannels, but atomic ion fluxes were undetectable under the same conditions (14). This anomaly in ionic and molecule permeation has also been described for other connexins, CALHM-1 and pannexin-1 channels (15, 16, 19). A proposed explanation was that molecule permeation occurred during brief and experimentally unresolved opening of the channels (14). In this study, we demonstrated that molecules can permeate in the absence of ionic conductance, suggesting that molecules and atomic ions are transported by different channel conformations or dynamics. Consistent with this notion, DAPI did not affect macroscopic or unitary ionic currents of Cx26 hemichannels as would be expected for molecules that occlude or bind within the pore at the same time that ions are diffusing through, as it has been shown using nanopores (36, 37). In a clear demonstration of the disconnect between ion and molecular permeability, we showed that pathological human mutations in the NT domain that were cataloged as loss-of-function mutants because of the absence of ionic currents, still retained the capability to transport molecules with similar kinetic parameters to those observed for wild-type channels. These findings indicate that molecules and ion permeation can be uncoupled.

The notion that ion and molecule permeation can be uncoupled is further supported by our finding that DAPI transport thorough Cx26 hemichannels is enabled at negative membrane potentials, but not at positive potentials, where the ion fluxes are substantial (Fig. 5B). This is consistent with a previous report showing that permeation at positive potentials is restricted to ions in Cx46 hemichannels (38). The differential voltage regulation of atomic ion permeation and molecule transport could be explained by hemichannels operating in two modes, a molecule-specific transport mode at negative potentials, and in a classical ion channel mode at positive voltages (see the proposed model in Fig. 8).

**Figure 8.**
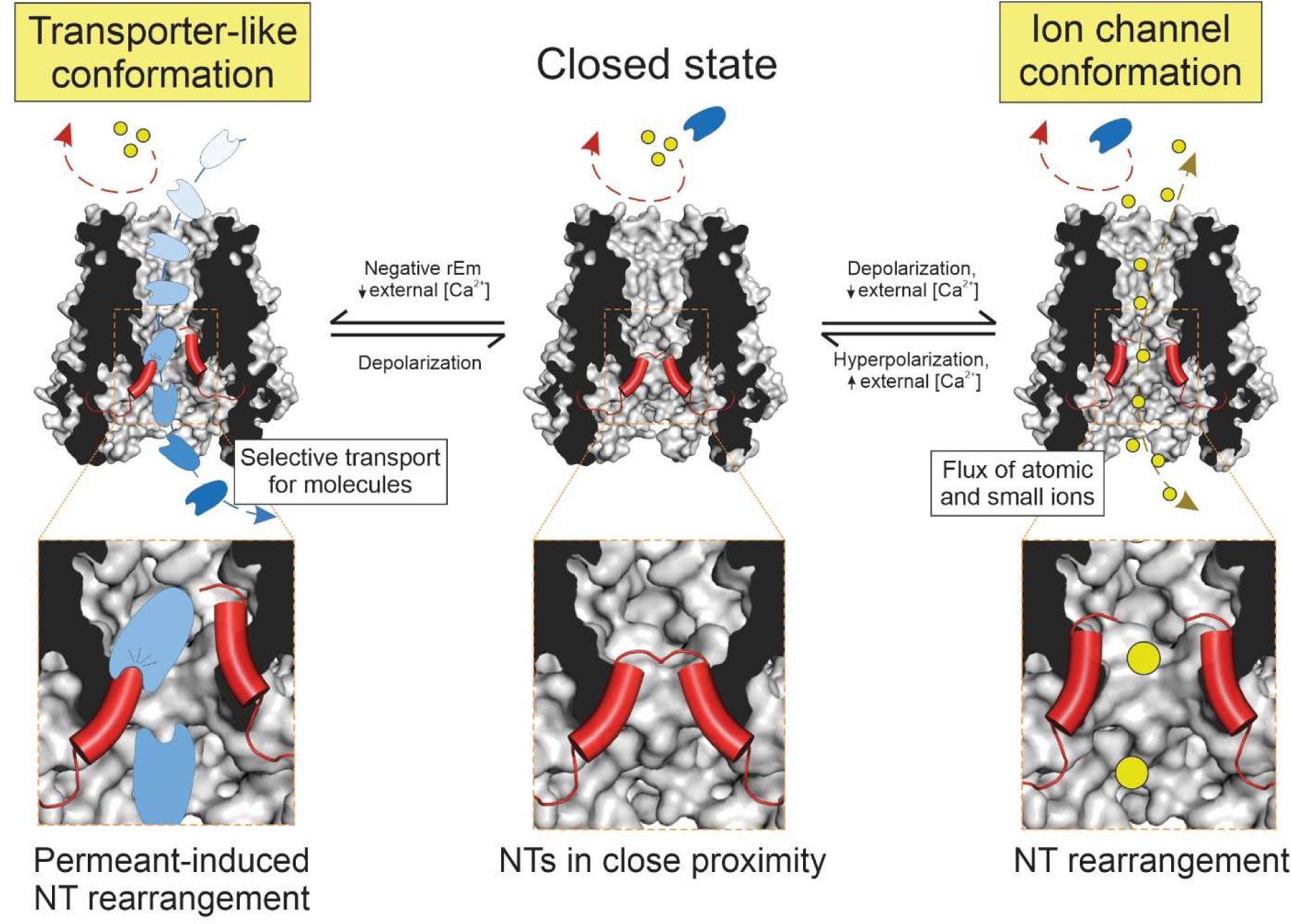
A proposed mechanism for the independent dual function of connexin hemichannels as molecule transporters and ion channels. In the closed state, hemichannels are not permeable to neither atomic ions or small molecules. At negative membrane potentials, a transporter-like conformation is favored, in which the interaction between permeants and specific residues lining the pore finely tune permeation properties, including transport kinetics and selectivity. We hypothesize that the interaction between permeants and the NT domain (shown in red), trigger a fast and short-term NT rearrangement that allows molecule translocation with no or negligible atomic ion leak. Notably, this mechanism is not disabled by physiological extracellular Ca^2+^ concentrations at which ionic conductance is negligible. Depolarization of the plasma membrane promotes the transition from a state non-conductive to ions to an open state that allows flux of atomic ions (e.g. K^+^, Na^+^, Ca^2+^, Cl^-^, represented as yellow circles) but not necessarily permeation of large molecules (represented in light blue). A low external Ca^2+^ concentration may differentially favor the transition to either the transporter-like state or the ion channel state. Under these conditions, ions can passively move through the pore following their electrochemical gradient, likely because of an allosteric coupling between the Ca^2+^-binding site and NT domain that keeps the gate open.

While our data suggest that the channels can favor permeation of either ions or molecules, we cannot rule out that, under specific conditions, a full open state can still mediate permeation of both at the same time, as is assumed for wide pores with large ionic conductance. In this work, we provide evidence that a transition to this expected functionality is not necessarily induced by voltage or low extracellular Ca^2+^. First, the fact that positive potentials abolish DAPI permeation suggests that voltage-induced activation allows the transition to an open state that is permeable to ions but not molecules as previously suggested (38). Second, we found that in the presence of 1 mM extracellular Ca^2+^ Cx26 hemichannel currents were reduced by ∼95%, but under the same conditions, DAPI permeation was reduced by only 50%, without effect on DAPI Km (Supplementary Fig. 14). Interestingly, a similar effect was recently observed in CALHM1 channels using a different cationic dye, YO-PRO-1 (19). Consistent with this, ionic conductance and ethidium transport are differentially affected by extracellular Ca^2+^ or Mg^2+^ in Cx30 hemichannels (39). Altogether, these data indicate that the magnitude of molecule transport does not correlate with the channel open probability reported by ionic conductance. We view it more likely that low Ca^2+^ can favor two states with preferential permeation to ions or to molecules but not to both simultaneously through the same pore configuration. Based on our findings, we hypothesize that extracellular Ca^2+^ might serve as a physiological modulator to maintain hemichannels closed to atomic ions preventing changes in resting membrane potential or the loss of electrochemical gradient, without disabling the mechanism for molecule transport.

### The NT domain is a flexible domain with critical roles in gating and molecule translocation

Structural studies have shown that the NT domain of Cx26 and Cx46/50 is folded into the pore, often parallel to the transmembrane segments, against the pore wall, as for other large-pore channels including innexins, pannexin, CALHM, and LRRC8 channels (6, 7, 40–46). However cryo-EM structures of Cx30.1, and Cx43 also show the NT domain separated from the pore wall and lying parallel to the axis of the plasma membrane, thereby narrowing the pore (9, 10). These and other data suggest that the NT of connexin channels is a highly flexible intra-pore domain, and therefore might serve as a gate for permeants (47). Here, by using extracellular Ca^2+^ as a connexin gating modulator, we experimentally demonstrated rearrangements of the NT domain in a state-dependent manner. The NTs of adjacent subunits come into proximity when the channel is closed by extracellular Ca^2+^. Notably, Ca^2+^ binding sites are localized in the extracellular part of the pore (24, 48); this also implies that there is an allosteric coupling between the Ca^2+^ sensing domain at the extracellular side and the NT at the intracellular side of the pore.

We propose here that movements of individual NTs are necessary to translocate small molecules across the pore with negligible fluxes of atomic ions. We support this notion with MD simulations showing that molecules directly interact with one or two NTs, triggering fast and short-term rearrangements. Experimental data support these interactions, showing that immobilization of the ends of the NT domains via disulfide bridges prevent molecule transport. Importantly, individual NT rearrangement has been already observed in structural studies for Cx43 (11).

Further studies must be carried out to elucidate how a large-pore channel may transport molecules with an undetectable flux of atomic ions. Based on our data we hypothesize that the dynamics of the pore with a permeant present and the proper interaction between the permeant with the NT domain in a closed state can trigger a fast and short-term movement of the NT domain that adjust to the closed state immediately after the permeant leaves the NT region (i.e., a facilitated diffusion-like mechanism in which interaction of molecules with the NT induce their own translocation). Alternatively, a second gate/barrier specific for atomic ions could be present in our predicted conformation that allow molecule but not ion permeation. Either scenario would satisfy the minimal requirement for a transporter-like process, in which substrate binding induces a conformational change that shifts the barrier to substrate movement from one side of the substrate to the other (49).

### Implications for the current understanding of permselectivity properties in gap junction channels

Early reports showed that permeation of small ions (e.g., TEA^+^) through GJCs follows a linear relationship with GJC conductance (12). However, this correlation is not observed with fluorescent dyes of higher molecular weight (50), indicating that molecular permeability is potentially regulated in a different way than ionic permeation, as suggested here for hemichannels. Notably, early studies suggest that just a small fraction of gap junction channels is responsible for electrical coupling (51). Whether the remaining “closed” GJCs contribute to molecule transport independent of ionic conduction is a question that remains to be addressed.

The observation that selectivity on GJCs is isoform-specific and that the size of permeants does not necessarily correlate with the single channel conductance has suggested that the major determinants for permselectivity through GJCs could rely on interactions inside the pore more than the pore width per se (52, 53). Recent structures of Cx43 obtained by Cryo-EM and single particle analysis show that the overall architecture of hemichannels and gap junctions is similar. Moreover, the electrostatic properties of the pore are comparable (10, 28), suggesting that the residues lining the pore are not significantly affected when two hemichannels dock to form one GJC. We hypothesize that interactions between permeants and the residues lining the pore of undocked hemichannels might be conserved in GJCs, though one could expect that docking of hemichannels has, to some extent, an allosteric effect on the dynamics of the NT domains and how they interact with molecular permeants.

The findings from this work have critical implications for the physiological and pathological role of connexin hemichannels. For instance, electrostatic and hydrophobic interactions with the NT are likely the major determinants of selectivity and permeation kinetics for molecules that fit within the pore. Secondly, permeation of molecules can be uncoupled from atomic ion flux and rely on a “transporter-like” state impermeable to atomic ions. This state is favored at negative resting potentials and not disabled at physiological millimolar Ca^2+^ concentrations, in which connexin-dependent atomic ion flux is almost negligible. Connexin hemichannels might have distinguishable roles as ion channels and molecule transporters in health and disease. This implies that the mechanisms by which connexin human mutations produce diseases, specifically regarding molecular signaling through hemichannels and perhaps GJCs, should be re-evaluated.

## MATERIALS AND METHODS

### Molecular biology

cDNAs for human Cx26 and Cx30 were synthesized and subcloned by Epoch Life Science into pGEM-HA (Promega), or pcDNA 3.1^+^ (Thermofisher Scientific, USA) vectors for expression in Xenopus oocytes and HeLa cells, respectively. The cDNAs were transcribed *in vitro* to cRNAs using the HiScribe® T7 Arca mRNA Kit (New England BioLabs). Single mutations in Cx26 (W3A, Cx26 +3A, Cx26B, Cx26B +2C, D2N, N14Y and S17F) and Cx30 (T5M) were produced using the QuickChange II Site-Directed Mutagenesis Kit (Agilent Technologies) and confirmed by DNA sequencing.

### Xenopus oocytes for heterologous expression of connexin hemichannels

Oocytes were collected from female *Xenopus laevis* (Xenopus 1 Corp, Dexter, MI) as we previously described (20) according to the protocol approved by the Institutional Animal Care and Use Committee (IACUC) at University of California Davis and conforming to the National Institutes of Health Guide for the Care and Use of Laboratory Animals. Briefly, an ovarian lobe was digested using an OR-2 solution (composition in mM: 82.5 NaCl, 2.5 KCl, 1 MgCl_2_, 5 HEPES, adjusted to pH = 7.6) containing 0.6 mg/mL collagenase type-IA. Defolliculated oocytes (stage IV-V) were individually microinjected with connexin cRNAs (20-40 ng) plus a Cx38 antisense oligonucleotide (20 ng) using the Nanoliter 2020 Injector system (World Precision Instruments, Sarasota, FL, USA). Following microinjection, oocytes were stored in individual well plates containing ND96 solution (composition in mM: 96 NaCl, 2 KCl, 1 MgCl_2_, 1.8 CaCl_2_, 5 HEPES, adjusted to pH = 7.4) supplemented with streptomycin (50 µg/mL) plus penicillin (50 Units/mL). Oocytes were stored at 16 °C for 48 h before experiments.

### Oocyte preparation for fluorescence-based dye uptake assays

Frog oocytes were made translucent using an optimized protocol recently described in detail by us (19, 20). In brief, after a centrifugation process to render the animal pole of oocytes translucent, we microinjected the translucent zone of each oocyte with 50 nL of a solution containing DNA extracted from salmon testes (0.5 mg/mL) along with a DNA fluorescent reporter (100 µM DAPI dilactate or 100 µM YO-PRO-1). The choice of the fluorescent reporter was based on compatibility with the excitation/emission wavelengths of the dye used for permeation assays. The co-injection of a fluorescent reporter with salmon DNA allowed us to adjust the focal plane in the injected DNA, enhancing the accuracy and sensitivity of the dye uptake assay. The injected DNA also provided an excess of binding sites, as previously characterized (19). Following microinjection, oocytes were stored in ND96 solution at 16 °C for a minimum of 90 min to ensure equilibration before conducting any experiments.

### TEVC/Dye uptake assay

Dye uptake was evaluated simultaneously with the Two-Electrode Voltage-Clamp (TEVC) configuration as previously described (19, 20). Translucent oocytes were positioned in a perfusion chamber on the stage of an inverted epifluorescence microscope (Nikon Eclipse Ti), with the translucent side of the oocyte facing a 20X objective. Oocytes were impaled at the vegetal pole using two electrodes filled with 3 M KCl (resistance: 0.2 - 2 MΩ). Dye uptake was recorded using a photomultiplier tube (Thorlabs, PMT1002) connected to a data acquisition system (Digidata 1440A, Molecular Devices, USA). Both dye uptake and electrophysiological recordings were acquired at a 2 kHz sample rate and analyzed using pClamp 10 software (Molecular Devices, USA).

The uptake of ethidium (314 Da) or DAPI (279 Da) was assessed at the resting membrane potential unless otherwise specified. Oocytes were allowed to equilibrate for 5-10 min after microelectrode impalement to recover. Only oocytes with a stable resting membrane potential were used for experiments. The values corresponding for the resting membrane potential for each experimental condition are shown in the Supplementary Table 1. Non-injected oocytes or oocytes injected with antisense alone were used as controls to measure batch-dependent endogenous dye uptake. Only oocyte batches with low or negligible background uptake were included in the analysis. Dye uptake measurements were conducted at room temperature (20-22 °C) in the absence of light. A Ringer’s solution (composition in mM: 115 NaCl, 2 KCl, 1 CaCl_2_, 5 HEPES, pH adjusted to 7.4) was used for dye uptake assessments, unless otherwise specified.

For measurements under voltage-clamp conditions, recording of dye uptake started 2-3 min after the voltage was clamped to a specific membrane potential. Dye uptake recordings in voltage-clamped oocytes were conducted for a duration of 10 min for each voltage tested. Since oocytes typically do not maintain steady currents after extended or repeated voltage clamping at extreme potentials, no more than 3-5 voltages were tested per oocyte.

### Determination of kinetic parameters of molecule permeation

The rates of dye uptake were obtained by calculating the slope of dye uptake recorded over time. Dye uptake along the time is expressed as arbitrary units (A.U.) and dye uptake rates are expressed as arbitrary units per second (A.U./s). To calculate the maximum transport rate (Vmax) and the apparent affinity for molecule transport (Km), curves were fit to a Michaelis-Menten equation using Origin 9 software (OriginLab Corporation, Northampton, MA, USA).

### Electrophysiology

For TEVC, two pulled borosilicate glass micropipettes were filled with 3 M KCl resulting in resistances ranging from 0.2 to 2 MΩ. Electrodes for voltage recordings and current injection were mounted on separate micromanipulators (Burleigh PCS-5000, USA) on the stage of a Nikon Eclipse Ti Microscope. The electrical signal was amplified using an Oocyte Clamp Amplifier (OC-725C, Warner Instrument Corp., USA) and digitized by a data acquisition system (Digidata 1440A, Molecular Devices, USA). Data were sampled at 2 kHz, acquired at room temperature (20-22 °C), and analyzed using pClamp 10 software (Molecular Devices, USA). TEVC was performed using Ringer’s solution (composition in mM: 115 NaCl, 2 KCl, 1 CaCl_2_, 5 HEPES, pH adjusted to 7.40) unless otherwise specified. Voltage-dependence curves were obtained by analyzing the magnitude of tail currents evoked by depolarizing pulses (-80 mV to +80 mV, holding potential: -80 mV, duration of depolarization pulse: 40 s), as previously described by Lopez et al. (48). To study connexin hemichannel opening by low Ca^2+^, oocytes were superfused (∼0.75 mL/min) with a Ringer’s solution containing 0.2 mM CaCl_2_. Current-voltage relationships were obtained by analyzing the magnitude of the ionic currents recorded during voltage step pulses (-60 mV to +60 mV, holding potential: -30 mV, duration of depolarization pulse: 0.5 s). To obtain Ca^2+^ concentration-response curves, oocytes were clamped at -40 mV and superfused (∼0.75 mL/min) with a Ringer’s solution containing 0.01 mM CaCl_2_ to increase the relative open probability of hemichannels. After currents stabilized, the magnitude of ionic current inhibition at each Ca^2+^ concentration was calculated and used for analysis. The Ca^2+^ concentration-response relationship was fitted to the Hill equation using Origin 9 software (OriginLab Corporation, Northampton, MA, USA).

Single-channel recordings of Cx26 hemichannels in Xenopus oocytes were performed following previously established methods with minor adjusments (54, 55). Vitelline layer-free oocytes were placed in a chamber filled with a patch solution (composition in mM: 140 KCl, 1.8 CaCl_2_, 1 MgCl_2_, 5 HEPES, pH adjusted to 7.40). Glass capillaries were pulled in a horizontal puller (Sutter Instrument Co.) and fire-polished using a Micro-forge (MF-200, World Precision Instruments) to obtain a resistance ranging from 1 MΩ to 8 MΩ when filled with the patch solution. To investigate the effect of molecular permeants on single-channel properties, we included 200 µM DAPI dilactate in the patch pipette. Patch clamp experiments were conducted using a Multiclamp 700B amplifier (Axon Instruments, Molecular Devices, CA, US). Data were digitized at 5 kHz using a 16-bit A/D converter data acquisition system (Axon Digidata 1550B, Axon Instruments, Molecular Devices, CA, US), and recorded using Clampex 11 acquisition software (Axon Instruments, Molecular Devices, CA, US). Single-channel activity was assessed in the cell-attached configuration. To calculate the command voltage needed to obtain a specific membrane potential at the patch site (that is, the membrane potential that the recorded channels see), we used the equation E_patch_ = E_m_ -V_Cmd_, where E_patch_ represents the membrane potential at the patch, E_m_ is the resting membrane potential of the oocyte, and V_Cmd_ is the command voltage applied in the patch pipette. All potentials described in this manuscript correspond to E_patch_.

### Molecular dynamics simulations

#### Atomistic model preparation

We utilized a previously established atomic model of the Cx26 hemichannel (21), which recapitulates the experimental single-channel permeability of cyclic-AMP (22). In this model, acetylated N-termini and methylated C-termini capping were used for the terminal amino acids of each segment. According to the crystal structure of the Cx26 junctional channel (Protein Data Bank: 2ZW3) (6), we kept three disulfide bonds per protomer (C53-C180, C64-C169, and C60-C174), resulting in a total of 18 disulfide bonds. This model was embedded within a solvated POPC bilayer, in the presence of ions and TIP3P water molecules. Two systems were simulated here, one of which contained a single molecule of ethidium, while the other incorporated a molecule of 4’,6-diamidino-2-phenylindole (DAPI). To maintain a physiological ionic environment and neutralize the protein’s charge, we introduced K^+^ and Cl^-^ ions into the system.

#### Force fields

We employed the CHARMM36 force field for various components of our system, including the protein (56, 57) 1-palmitoyl-2-oleoylphosphatidylcholine (POPC) lipids (58), as well as KCl and TIP3P water molecules (59). In addition, we optimized CHARMM General Force Field (CGenFF) (60) force field parameters (the atomic charge and dihedral angles) for DAPI and ethidium using FFParam (61).

#### Equilibrium and production simulation protocols

The construction and equilibration of the system were carried out in accordance with a step-by-step protocol provided by the Membrane Builder tool, accessible on the CHARMM-GUI website (62, 63). Briefly, the system underwent a rigorous energy minimization process and sequential equilibration in both NVT (constant Number of particles, Volume, and Temperature) and NPT (constant Number of particles, Pressure, and Temperature) ensembles, with the application of multiple steps of protein backbone and lipid headgroup positional restraints. Molecular dynamics simulation engine AMBER20 (64) was used throughout this study. Temperature control was maintained at 303.15 K using a Langevin thermostat (65, 66), and pressure was regulated at 1 atm through Monte Carlo barostat pressure control (67, 68). A time step of 2 fs was employed for all the simulations. The cutoff distances for van der Waals interactions and short-range electrostatic interactions were set at 12 Å, with a force switch occurring at 10 Å. For long-range electrostatic interactions, we utilized the particle mesh Ewald algorithm (69).

#### Voltage simulations

After a 200 ns unbiased simulation, -200 mV voltage was applied across the simulation box to drive the inward flux of dye molecules, in the same direction as the dye uptake experiments driven by concentration gradient. For each dye molecule, three replicas of 500 ns were carried out under a -200 mV voltage using AMBER with the same simulation protocols as above. Trajectories of the system were saved at regular intervals of 100 ps.

### Biotinylation and Western blotting

Oocytes expressing wild-type or mutant connexins were incubated in HEN solution (composition in mM: 250 HEPES, 1 EDTA, and 0.1 neocuproine, adjusted to pH = 7.7) containing 0.5 mg/mL EZ-Link^TM^ Sulfo-NHS-SS-Biotin (ThermoFisher Scientific, USA) for 20 minutes at 4° C. Subsequently, oocytes were washed with PBS containing 15 mM glycine (pH = 7.4) for 15 minutes. Oocytes were then homogenized in HEN solution in the presence of protease inhibitors (ThermoFisher Scientific, USA) and 1% Triton X-100, followed by centrifugation for 10 minutes at ∼13,500 g and 4° C. Next, 10 µL from the supernatant (representing the total fraction) was preserved for total protein analysis, while the remaining supernatant was incubated with streptavidin agarose (ThermoFisher Scientific, USA) for 60 minutes at 4° C. Samples were then centrifuged for 2 minutes at 14,000 g, and the supernatant was discarded. The fraction containing biotinylated proteins was washed with HEN solution plus protease inhibitors. Then, samples were gently mixed with Laemmli buffer + β-mercaptoethanol (Sigma-Aldrich, St. Louis, MO, USA) to disrupt the biotin–streptavidin interaction and heated for 5 minutes at 80° C. Finally, total proteins and biotinylated proteins were separated by 12% SDS-PAGE and transferred onto a PVDF membrane (BioRad, Hercules, CA, USA). The Signal Enhancer HIKARI (Nacalai Tesque, INC, Japan) was used for incubating the primary antibody (anti-Cx30, ThermoFisher Scientific, Cat. # 71-2200, 1:1,000 or anti-Cx26, Millipore, Cat. # MABT198, 1:1,000) and secondary antibody (Anti Rabbit IgG, ThermoFisher Scientific, Cat. # 32460 or Anti Mouse IgG, ThermoFisher Scientific, Cat. # 31430, 1:10,000). SuperSignal® West Femto (ThermoFisher Scientific, USA) was used for protein band detection. Molecular mass was estimated using pre-stained markers (BioRad, Hercules, CA, USA).

For crosslinking experiments, oocytes expressing Cx26B and Cx26B +2C were incubated for 10 min in Ringer K^+^ (composition in mM: 117 KCl, 5 HEPES, pH adjusted to 7.4) with or without TbHO_2_ (2 mM) or CaCl_2_ (5 mM). Oocytes were homogenized in a buffer containing 20 mM Tris-HCl (pH 7.6), 100 mM NaCl, 1% Triton X-100 and protease inhibitors. Lysates were centrifuged at 10,000 g for 2 min. The supernatant was separated and mixed with Laemmli buffer. Samples were then separated in a 12% Mini-PROTEAN TGX Precast Gel (BioRad, Hercules, CA, USA) and transferred to a 0.2 μm PVDF membrane. The membrane was incubated overnight with an Anti-HA antibody in TTBS buffer and developed as mentioned above. For quantification, four independent replicas were analyzed by using ImageJ software.

### Dye uptake measurements in HeLa cells

HeLa cells knockout for Cx45 were seeded on black 96-well plates and transfected with EGFP alone or in combination with connexin-containing pcDNA 3.1^+^ plasmids (Cx26^WT^, Cx26^N14Y^, Cx30^WT^, or Cx30^T5M^). Cells were transfected using the JetPrime® Transfection Reagent (Genesee Scientific, CA) which exhibited robust transfection efficiency with HeLa cells. 24 h after transfection, media components were removed by washing the cells twice with pre-warmed external solution (composition in mM: 142 NaCl, 4 KCl, 1 CaCl_2_, 5 D-glucose, 10 HEPES) adjusted to pH=7.40 and 285 mOsm, equivalent to the media osmolarity. EGFP signal was initially measured using a multimode plate reader (Victor Nivo, Perkin Elmer; excitation filter: 480/30, emission filter: 530/30). Next, the solution was replaced with the pre-warmed external solution (described above) containing 30 µM DAPI dilactate or 3 µM ethidium bromide. Cells were incubated for 15 min at 37 °C. Then, the total dye uptake was evaluated in the same plate reader [excitation filters: 355/40 (DAPI), 530/30 (ethidium), emission filters: 460/30 (DAPI), 580/20 (ethidium)]. Cell health was confirmed before and after the assay by optical visualization. Background signals were subtracted, and EGFP signal was used for data normalization. All measurements were performed at 37 °C.

### cAMP measurement

24 h after cRNA injection, oocytes were transferred to well-plates containing ND96 solution supplemented with 100 µM 2’5’-Dideoxyadenosine (DDA) to inhibit the endogenous production of cAMP. 24 h later, oocytes were transferred and incubated for 90 min in new wells containing ND96 solution supplemented with 100 µM DDA + 500 µM 3-Isobutyl-1-methylxanthine (IBMX) to inhibit phosphodiesterase activity. Then, oocytes were transferred to new plates containing a saline solution (composition in mM: 115 NaCl, 2 KCl, 5 HEPES, 0.5 EDTA, 0.5 EGTA, pH = 7.4) supplemented with 100 µM DDA + 500 µM IBMX and cAMP (0 -1,000 µM). Oocytes were incubated with cAMP at room temperature for 90 min with gentle shaking. After the cAMP incubation, oocytes were washed six times using a washing solution (composition in mM: 115 NaCl, 2 KCl, 5 HEPES, 10 CaCl_2_, 10 MgCl_2_, 0.2 LaCl_3_, 0.1 DDA, and 0.5 IBMX, pH = 7.4). Finally, oocytes were homogenized in 0.1M HCl and centrifuged at ∼19,000 g for 15 min, 4 °C. cAMP levels from the lysates were measured using the Direct cAMP ELISA Kit (Enzo Life Sciences) following the manufacturer’s instructions.

### Reagents

MS-222 was obtained from Syndel (Ferndale, WA). Collagenase type IA, Ficoll® PM 400, salmon DNA, HEPES, DDA, IBMX, glycine, neocuproine, ethidium bromide, TBHO_2_, and cadmium chloride, were purchased from Sigma-Aldrich (St. Louis, MO, USA). Penicillin-streptomycin, DAPI dilactate and YO-PRO^TM^-1 Iodide was obtained from ThermoFisher Scientific (USA).

### Statistical analysis

Values are presented as mean ± SEM. Comparisons between groups were made using paired or unpaired Student’s t-test, one-way ANOVA plus Newman-Keuls post hoc test, or two-way ANOVA plus Tukeýs post hoc test, as appropriate. P < 0.05 was considered significant.

## ACKNOWLEDGMENTS

This work was supported by the National Institutes of Health/National Institute of General Medical Sciences (Grants R01-GM099490 to J.E. Contreras and Y. L. Luo, and R01-GM101950 to A.L. Harris and J.E. Contreras). Computational resources were provided via the Extreme Science and Engineering Discovery Environment (XSEDE) allocation TG-MCB160119, which is supported by NSF grant number ACI-154862. This work is dedicated to the memory of Mike Bennett.

The authors declare no competing financial interests.

## AUTHOR CONTRIBUTIONS

JEC and PSG designed the research. YL and PSG performed molecular biology. PSG, DK, CIF, and JMVC performed research. DK conducted ethidium and DAPI simulations. YCL and WJ prepared ethidium simulation system and conducted simulations. AB prepared Force Field. PSG, DK, CIF, and JMVC analyzed data. PSG, DK, ALH, YLL, and JEC wrote the manuscript. PSG, ALH, YLL and JEC edited the manuscript.

