## Supplementary Figures for "Large-pore connexin hemichannels function as molecule transporters independently of ion conduction"

### SUPPLEMENTARY DATA

### SUPPLEMENTARY FIGURES

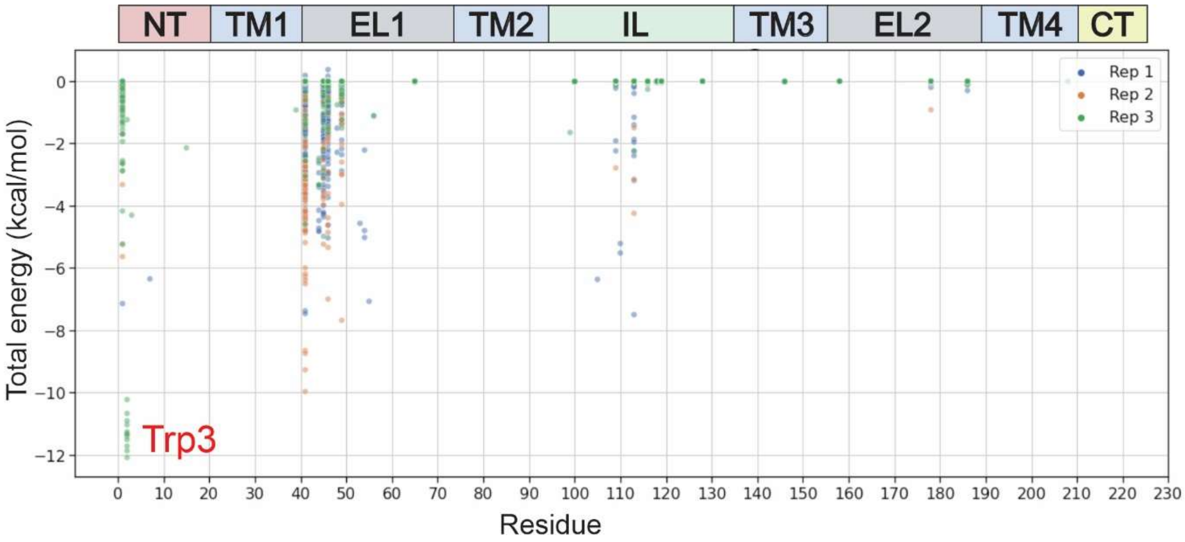


**Supplementary Fig. 1:** Energy interactions for ethidium in the pore of Cx26 hemichannels. An MMGB pairwise analysis was performed on 100 frames to calculate the interaction energies between ethidium and the residues lining the pore of Cx26 hemichannels (at −200 mV). The total energy refers to the overall energy contribution from van der Waals, electrostatic, polar and non-polar interactions.

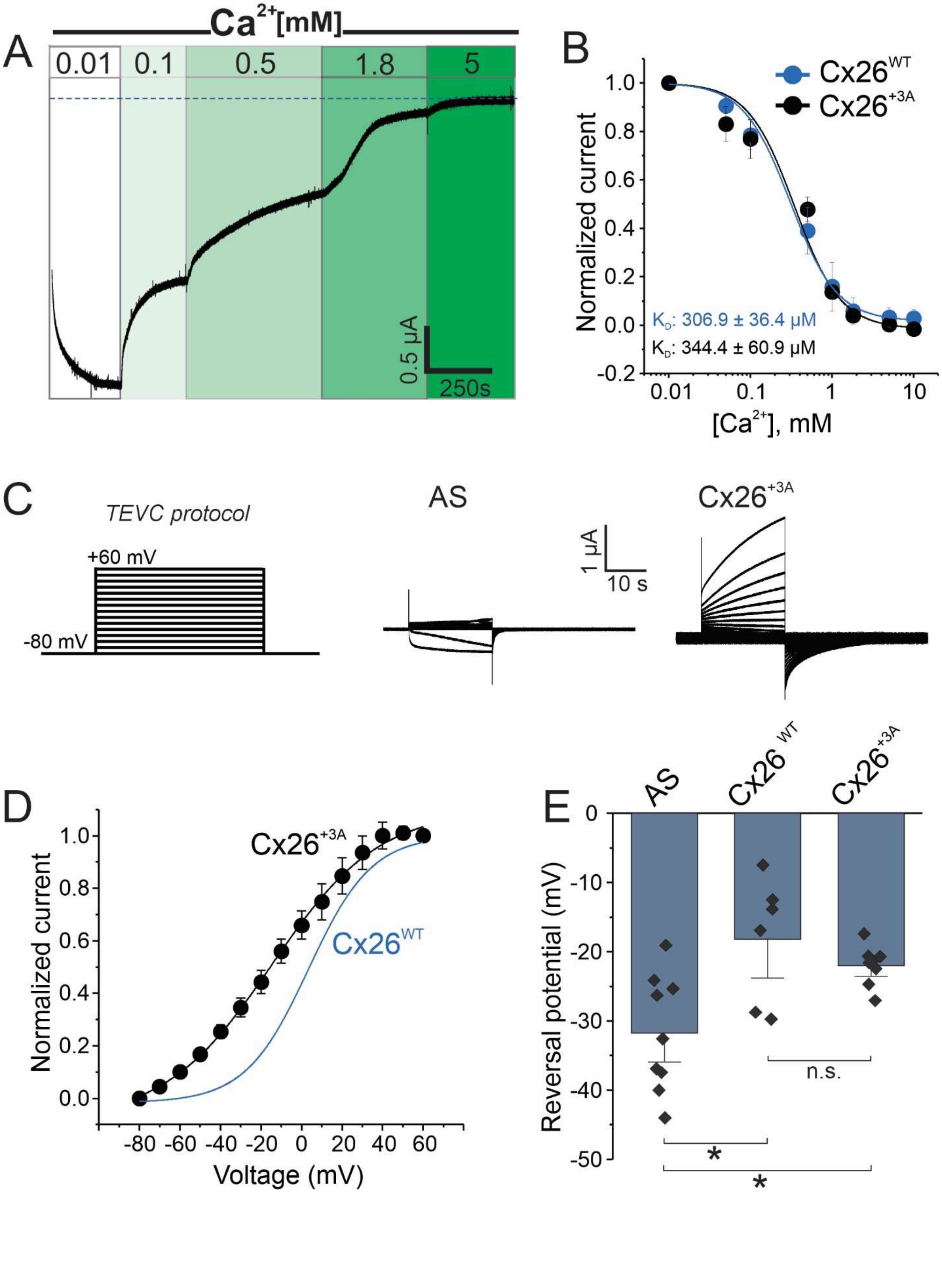

A
Ca2+[mM]
0.01
0.1
0.5
1.8
5
0.5 μA
250s
B
Cx26WT
Cx26+3A
Normalized current
KD: 306.9 ± 36.4 μM
KD: 344.4 ± 60.9 μM
[Ca2+], mM
C
TEVC protocol
+60 mV
-80 mV
AS
Cx26+3A
1 μA
10 s
D
Normalized current
Cx26+3A
Cx26WT
Voltage (mV)
E
AS
Cx26 WT
Cx26+3A
Reversal potential (mV)
n.s.
*
*

**Supplementary Fig. 2:** hCx26 +3A mutant displays biophysical properties of atomic ion permeation similar to WT hemichannels. Biophysical properties of Cx26 +3A hemichannels were evaluated by TEVC. **(A)** Representative trace of the ionic current recorded in oocytes expressing Cx26 WT. Oocyte was perfused with a Ringer solution containing 0.01 mM $Ca^{2+}$ (at holding potential -40 mV) to increase the relative open probability of the hemichannels. After ionic current stabilization, oocytes were perfused with increasing $Ca^{2+}$ concentrations, which reduced the relative open probability in a concentration-dependent manner. The blue dashed line indicates 0 μA. **(B)** Quantification of $Ca^{2+}$ sensitivity for Cx26 WT and Cx26 +3A following the experimental approach shown in **A**. Ionic current was normalized to the current values detected at 0.01 mM $Ca^{2+}$ (i.e., the maximum relative open probability). **(C)** TEVC protocol used to evaluate the voltage-dependent activation of Cx26 hemichannels. A voltage step of 40 s duration with a holding potential of -80 mV was used. The representative ionic current traces using this TEVC protocol with oocytes expressing Cx26 +3A are also shown. Oocytes injected with an antisense (AS) oligonucleotide to reduce the endogenous expression of Cx38 were used as control. **(D)** Current-voltage relationship for Cx26 +3A (black) and Cx26 WT (blue). **(E)** Reversal potentials recorded in oocytes expressing Cx26 WT, and Cx26 +3A. Note that reversal potential for Cx26 WT and Cx26 +3A are similar. *, $P < 0.05$ vs AS, by one-way ANOVA plus Newman-Keuls post hoc test. n.s.= non-significant. Error bars are SEM.

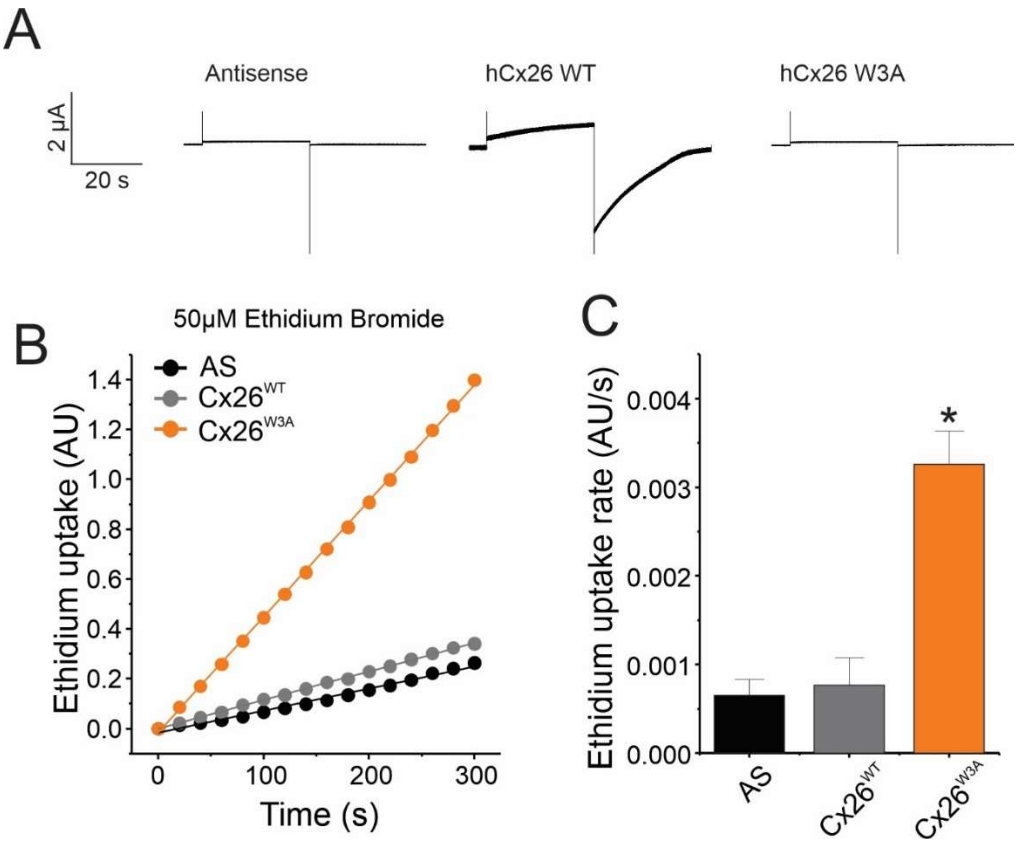


**Supplementary Fig. 3:** W3A mutation enables ethidium permeability and disables atomic ion conductance in Cx26 hemichannels. **(A)** Ionic currents were tested by TEVC in oocytes expressing Cx26 WT, or the mutant Cx26 W3A. Oocytes injected with an antisense (AS) oligonucleotide were used as control. Membrane potential was held at -80 mV to reduce the relative open probability of Cx26 hemichannels. Then, a 40 s depolarizing pulse (0 mV) was used to induce Cx26 hemichannel opening. After the depolarizing pulse, voltage was returned to the holding potential (-80 mV). **(B)** Representative time courses of ethidium uptake in oocytes incubated with 50 μM ethidium bromide. **(C)** Quantification of ethidium uptake rates taken from experiments shown in **B**. Note that W3A mutation enables ethidium permeability but disables ionic currents activated by voltage. *, $P < 0.05$ vs AS, by one-way ANOVA plus Newman-Keuls post hoc test. Error bars are SEM.

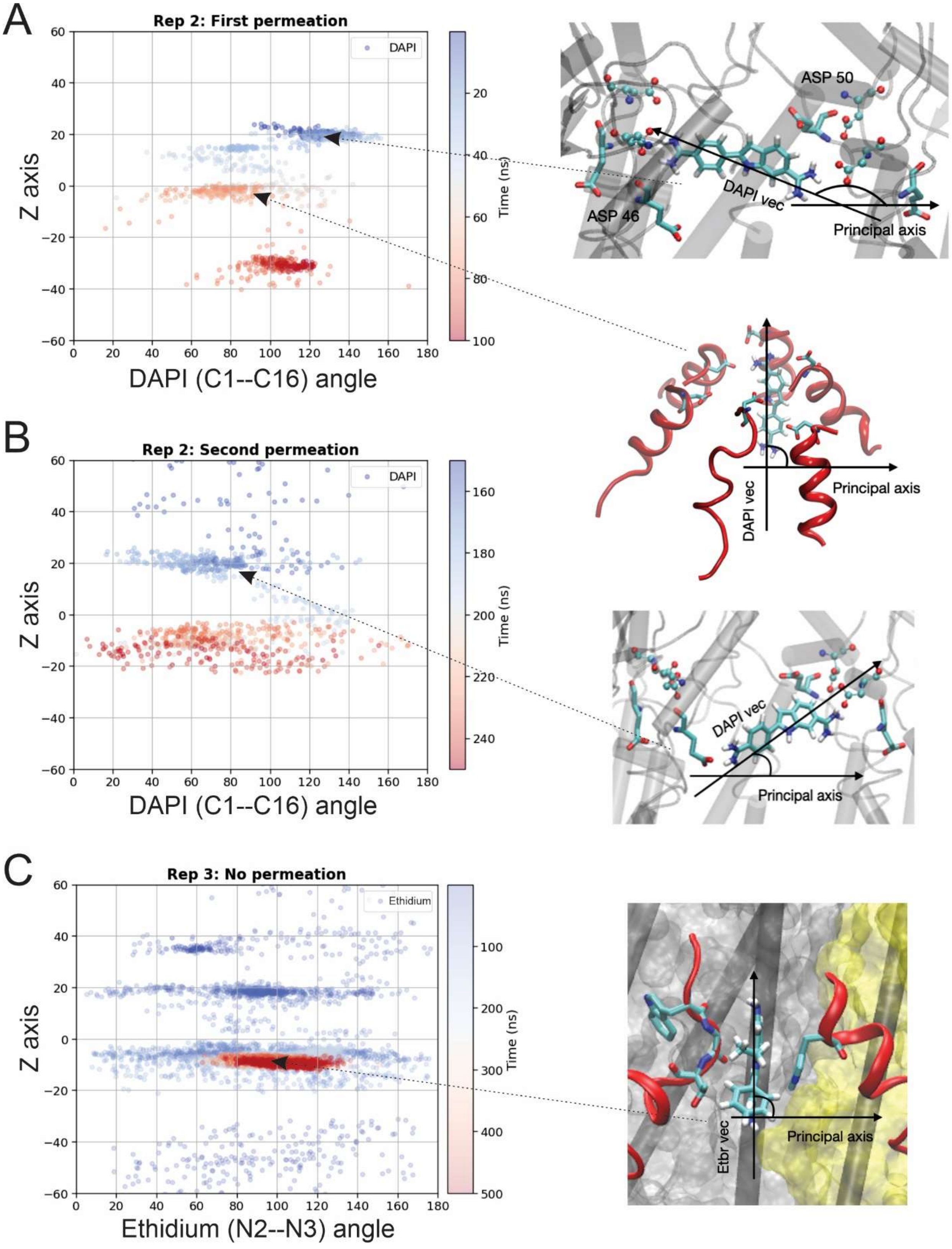

A
Rep 2: First permeation
DAPI
Z axis
DAPI (C1--C16) angle
Time (ns)
ASP 50
ASP 46
DAPI vec
Principal axis
B
Rep 2: Second permeation
DAPI
Z axis
DAPI (C1--C16) angle
Time (ns)
DAPI vec
Principal axis
DAPI vec
Principal axis
C
Rep 3: No permeation
Ethidium
Z axis
Ethidium (N2--N3) angle
Time (ns)
Etbr vec
Principal axis

**Supplementary Fig. 4:** Interactions of DAPI and ethidium with residues lining the pore contribute to molecule orientation inside the pore. MD simulations were performed using our Cx26 hemichannel model in the presence of DAPI **(A-B)** or ethidium **(C)** under -200 mV. The Z-axis orientation of molecules is determined relative to the center of mass of DAPI. The molecular angle is computed along the C1—C16 axis for DAPI and the N2—N3 axis for ethidium, in reference to the principal axis of lipid bilayer, represented along the X-axis. Panel **A** and Panel **B** depict the two distinct permeation events observed during 500 ns simulations in replica 2 (Rep 2, Fig. 3A). Panel **A** illustrates the first permeation occurring between 0 and 100 ns, while Panel **B** portrays the second permeation observed between 150 and 250 ns. Notably, both permeation events reveal distinct orientations within the extracellular and N-terminal (NT) region within the Cx26 pore. Specifically, within the NT region, DAPI exclusively assumes a vertical orientation, approximately 90 degrees relative to the lipid bilayer, though it is not perfectly vertical **(A)**. Conversely, within the extracellular region, DAPI molecules exhibit both horizontal and inclined orientations **(B)**. In contrast, ethidium does not exhibit permeation within the 500 ns simulation duration in replica 3 (Rep 3, Fig. 2A) **(C)**. Initially, during the production simulation, ethidium resided within the intracellular region, approximately between Z= -40 to -60 Å. Due to small size, ethidium demonstrates versatility in adopting multiple orientations within the extracellular region. However, within the NT region, ethidium establishes pi-pi stacking interactions with Trp3 in a vertical alignment, approximately 90 degrees relative to the bilayer, depicted by red dots. At the NT region, the chances of establishing pi-pi stacking are high in the vertical orientation at the NT region.

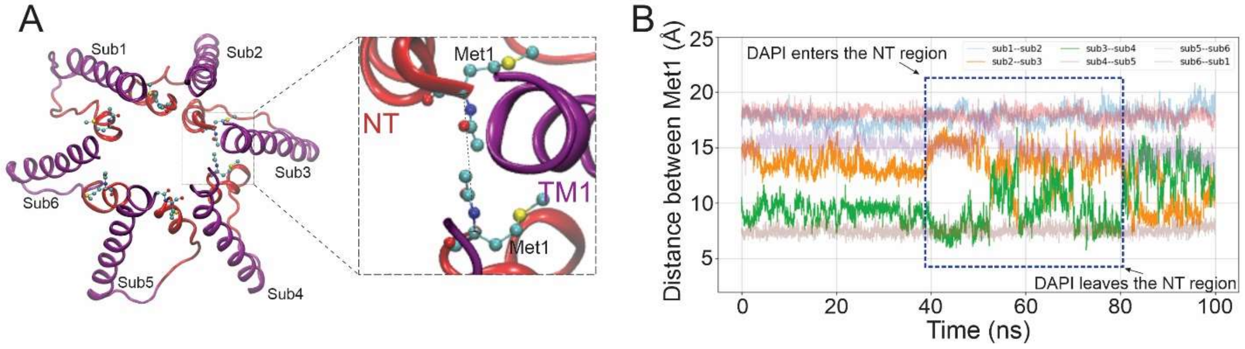


**Supplementary Fig. 5:** DAPI permeation induces NT rearrangements. MD simulations using an equilibrated Cx26 hemichannel in the presence of DAPI were used to determine NT dynamics during a permeation event. Voltage (-200 mV) was applied to simulate inward permeation and increase permeation events. **(A)** A cytosolic view of the Cx26 pore is represented in ribbon. Only TM1, (purple) and the NT domain (red) are shown for clarity. Met1 residues are represented in licorice with atom color code (red, oxygen; cyan, carbon; blue, nitrogen; yellow, sulfur; white, hydrogen). The inset highlights the distance between Met1 from adjacent subunits used for analysis in panel B. **(B)** Time course of the distances between Met1 residues from adjacent Cx26 subunits. The inset shows the period in which DAPI interacts with the NT during a single permeation event. Note that NT rearrangements occur when DAPI pass through the NT region.

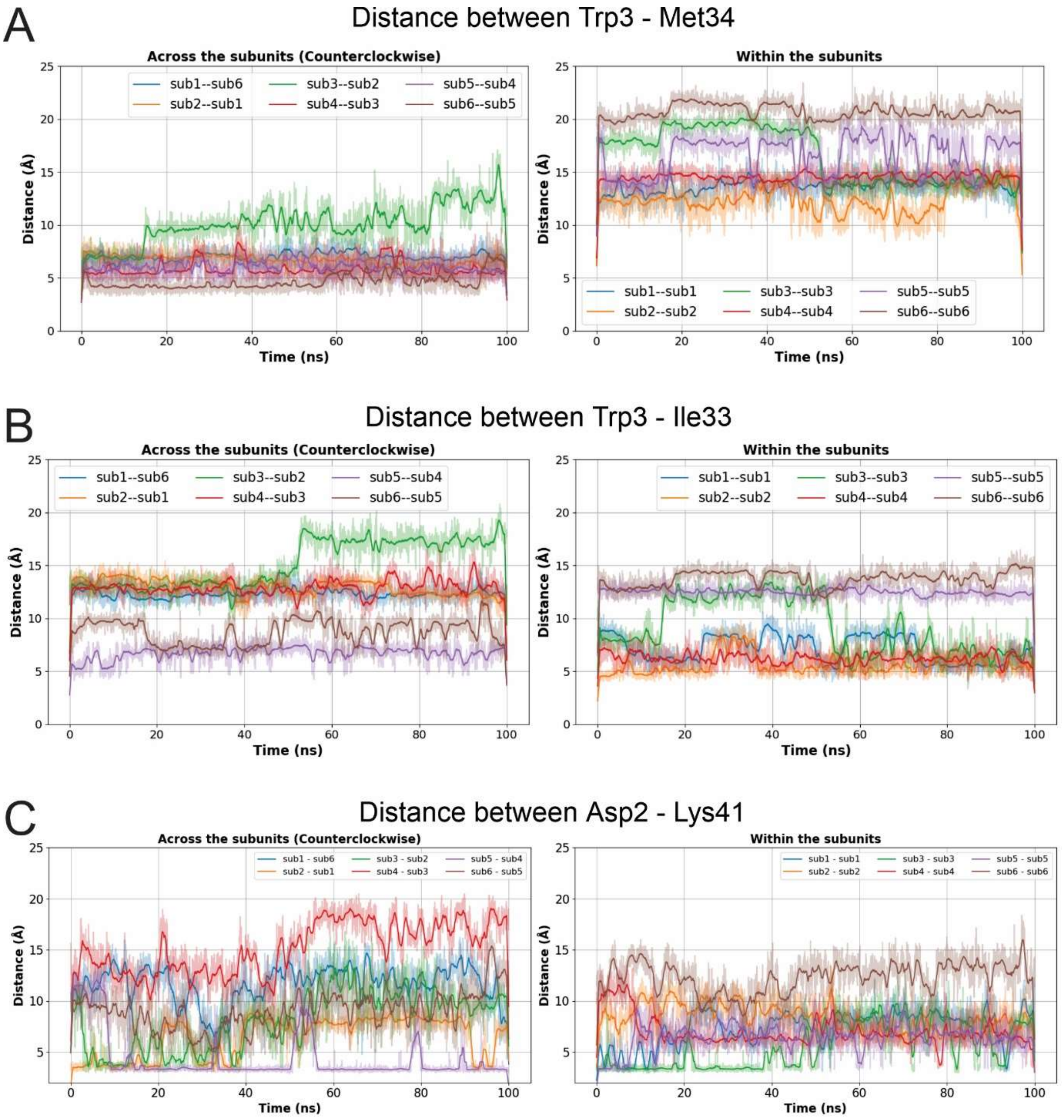


**Supplementary Fig. 6:** Interaction of DAPI with the NT is coupled with a disruption of interactions between the NT domain and TM1. MD simulations (under -200 mV) were used to determine NT dynamics in the Cx26 hemichannel during a permeation event. The distance between Trp3 - Met34 **(A)**, Trp3 - Ile33 **(B)**, and Asp2 – Lys41 **(C)** from adjacent subunits (left) or from the same subunits (right) is shown. The insets in each panel show the period in which DAPI interacts with the NT during a single permeation event.

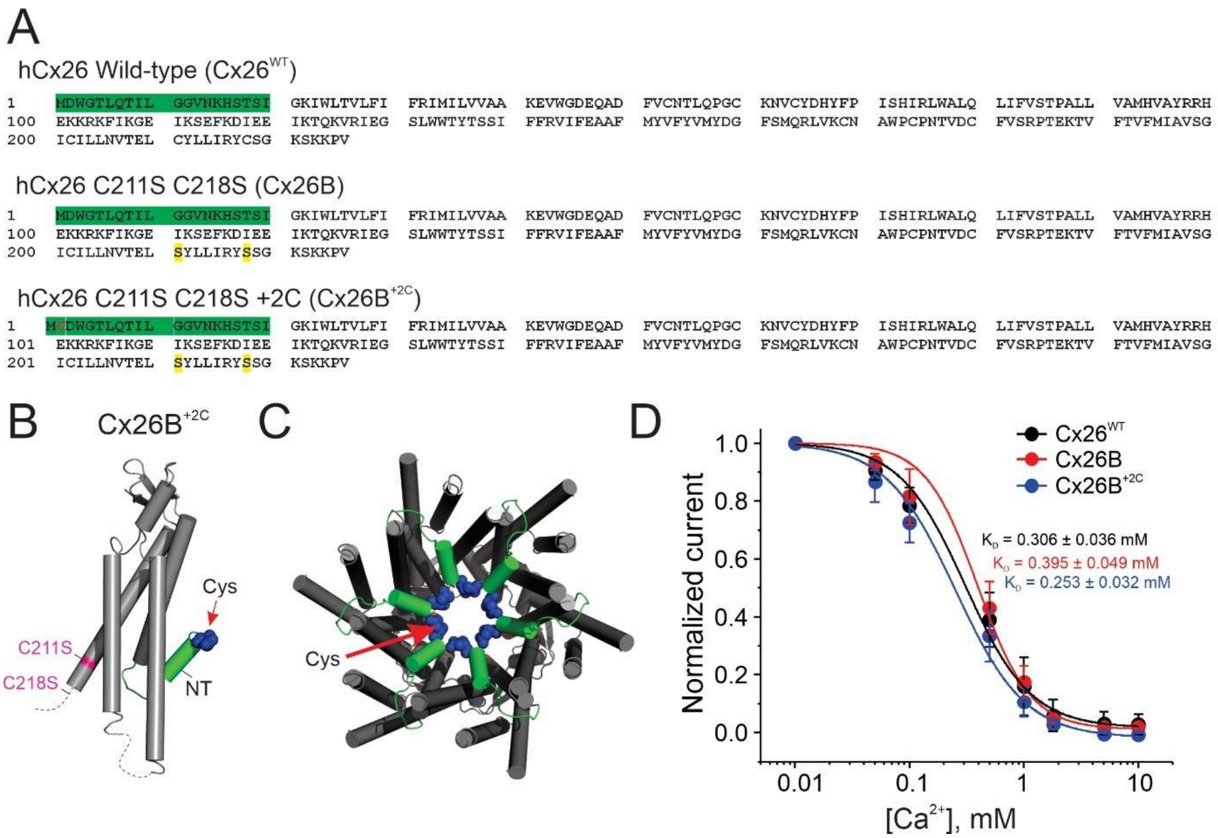


**Supplementary Fig. 7:** Incorporation of cysteine at position 2 in Cx26 leads to functional channels with similar $Ca^{2+}$ sensitivity. The addition of one cysteine at position 2 was used to evaluate crosslinking between NT domains. **(A)** Amino acid sequence of the human Cx26, Cx26B, and Cx26B +2C used in this work. The NT domain is highlighted in green. The incorporation of a cysteine at position 2 is shown in red, and the replacement of cysteines 211 and 218 for serine is highlighted in yellow. **(B)** Cx26 +2C monomer highlighting the cysteine (blue spheres) incorporated at position 2. **(C)** Bottom view of Cx26 hemichannel highlighting the cysteine residues incorporated in the NT domain lining the pore. **(D)** Ionic currents evaluated by TEVC in oocytes expressing Cx26 WT, Cx26B, and Cx26B +2C. Oocytes were perfused with Ringer solution with different concentrations of $Ca^{2+}$, at a holding potential -40 mV, as indicated in Supplementary Fig. 2A-B. Current was normalized to the maximum current obtained at 0.01 mM $Ca^{2+}$. Error bars are SEM.

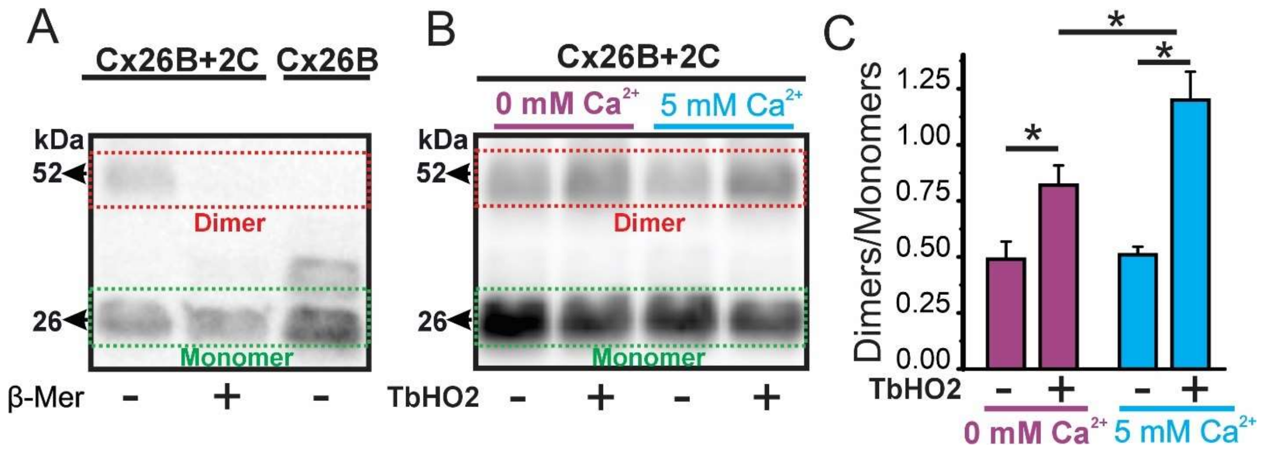


**Supplementary Fig. 8:** Extracellular $Ca^{2+}$ promotes disulfide bridge formation between Cx26B +2C subunits. **(A)** Representative western blot showing Cx26B and Cx26B+2C (~26 kDa each monomer) in absence or presence of β-mercaptoethanol (β-Mer). **(B)** Representative Western blot image showing Cx26B +2C treated with $TbHO_2$ for 10 min in a bath solution containing zero or 5 mM $Ca^{2+}$. **(C)** Quantification of dimer/monomer ratio for each condition showed in **B** (n = 4). *, $P < 0.05$, by two-way ANOVA plus Tukey´s post hoc test. Error bars are SEM.

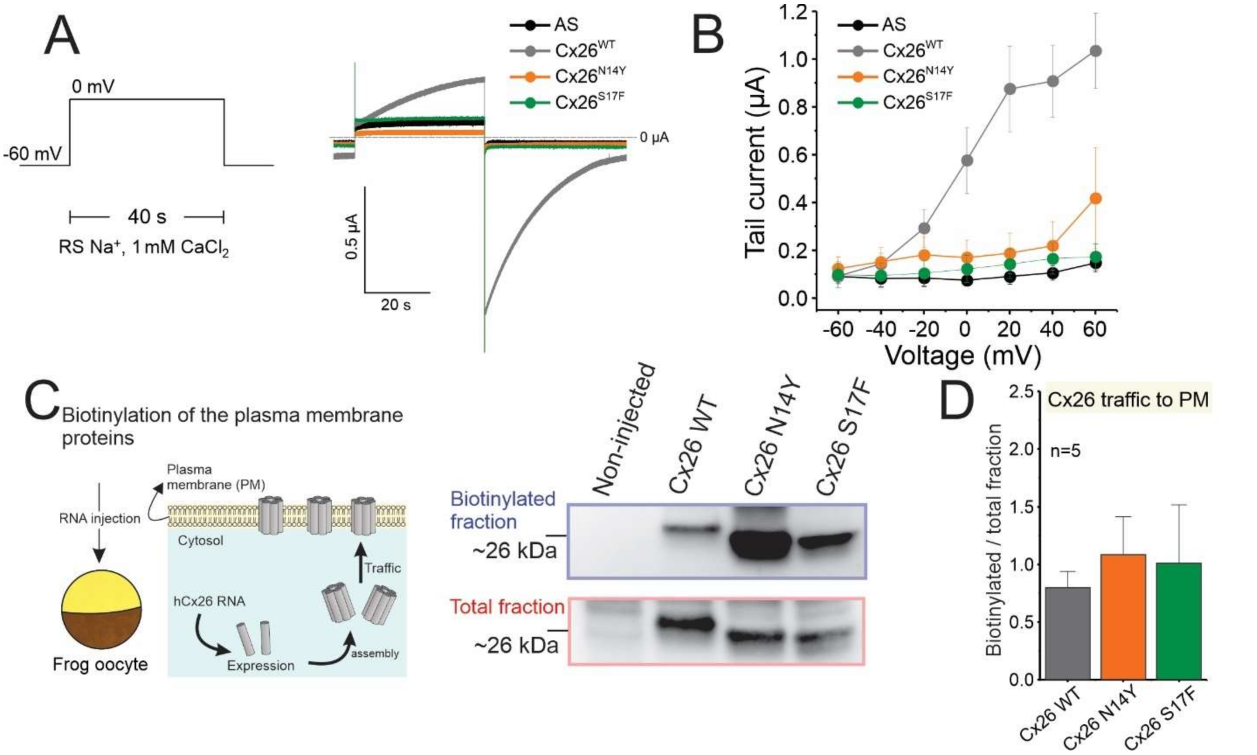


**Supplementary Fig. 9:** Human mutations in N14Y and S17F disable atomic ion permeation, but do not alter the traffic of Cx26 hemichannels to the plasma membrane when expressed in Xenopus oocytes. **(A)** Ionic currents were tested by TEVC in oocytes expressing Cx26 WT or the mutants. Oocytes injected with an antisense (AS) oligonucleotide were used as control. Oocytes were perfused with Ringer solution (RS) containing 1 mM $CaCl_2$. Membrane potential was held at -60 mV to reduce the relative open probability of Cx26 hemichannels. Then, a 40 s depolarizing pulse (0 mV) was used to induce Cx26 hemichannel opening. After the depolarizing pulse, voltage was returned to the holding potential (-60 mV) to visualize the tail currents (hemichannel deactivation). This protocol differs with the one shown in Fig. 7A-C since the hemichannel opening is induced by voltage instead of low extracellular $Ca^{2+}$. This second protocol corroborates that ionic conductance is disabled in these mutants. **(B)** Quantification of tail currents shown in A (n=6). **(C)** Scheme of the biotinylation approach. First, oocytes were injected with WT or mutant Cx26 RNA. After 2 days, oocytes were treated with biotin to label and separate plasma membrane proteins.

Antibodies targeting Cx26 were used to develop membranes (see Methods). Non-injected oocytes were used as a control. Both Cx26 mutants were found in the plasma membrane (i.e., in the biotinylated fraction, n=5). **(D)** Quantification of the Western blot bands suggest that mutations do not affect Cx26 traffic to the plasma membrane (PM). Statistic was performed using one-way ANOVA plus Newman-Keuls post hoc test. Error bars are SEM.

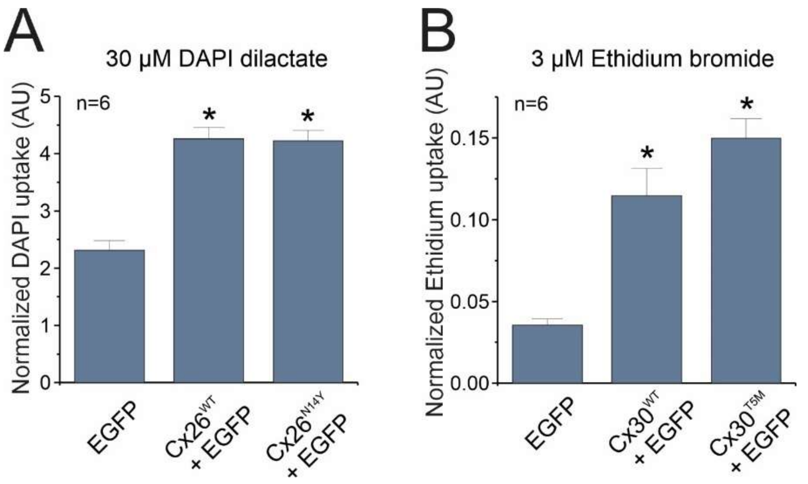


**Supplementary Fig. 10:** So-called loss-of-function pathological human mutations Cx26$^{N14Y}$ and Cx30$^{T5M}$ are permeable to fluorescent dyes when expressed in mammalian cells. HeLa cells deficient of connexins were transfected with EGFP alone (control group) or co-transfected with connexins (Cx26$^{WT}$ or Cx26$^{N14Y}$, in panel **(A)**, and Cx30$^{WT}$ or Cx30$^{T5M}$ in panel **(B)**. Cells were incubated 15 min at 37°C with an external solution containing 30 μM DAPI dilactate **(A)** or 3 μM ethidium bromide **(B)**. Dye uptake was evaluated in a fluorescent multimode plate reader (Victor Nivo, Perkin Elmer). EGFP signal was used to normalize data. *, $P < 0.05$ vs EGFP, by one-way ANOVA plus Newman-Keuls post hoc test. Error bars are SEM.

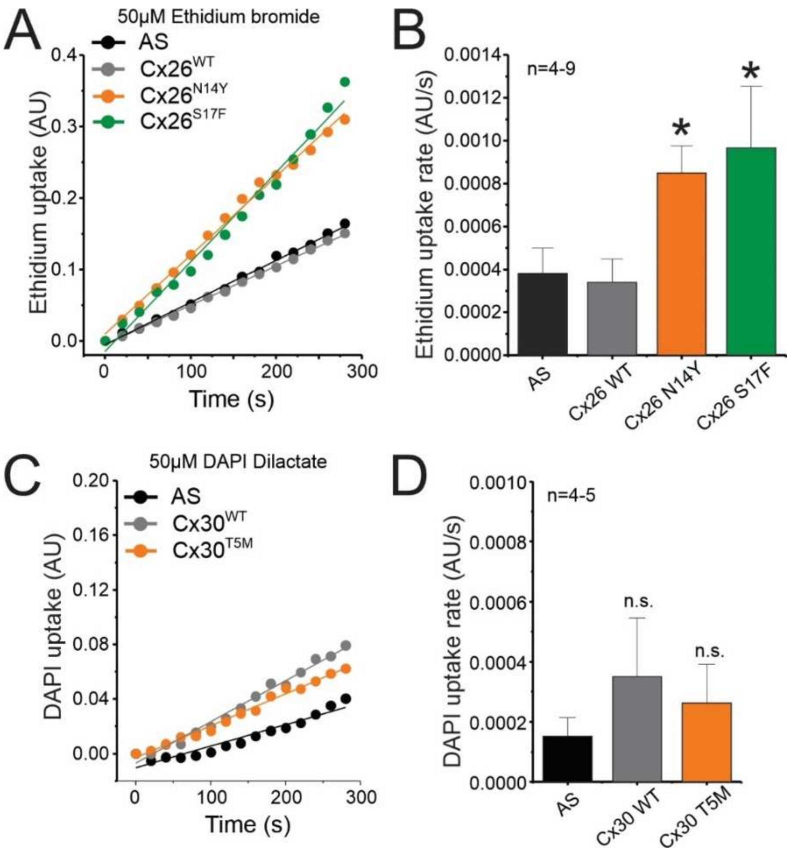


**Supplementary Fig. 11:** Human pathological mutations in the NT domain of Cx26 alter the selectivity to molecules. Molecular permeability studies were evaluated using the TEVC/dye uptake assay (see Methods). **(A)** Representative time courses of ethidium uptake in oocytes incubated with 50 µM ethidium bromide. Oocytes injected with an antisense (AS) oligonucleotide were used as a control to determine endogenous/unspecific ethidium transport. **(B)** Quantification of ethidium uptake rates from data shown in **A.** Note that both Cx26 mutants tested (N14Y and S17F) enabled permeation to ethidium. **(C)** Representative time courses of DAPI uptake in oocytes incubated with 50 µM DAPI dilactate. **(D)** Quantification of DAPI uptake rates from data shown in **C.** Note that the T5M mutation in Cx30 did not affect impermeability to DAPI. All measurements were done using 1 mM extracellular $Ca^{2+}$ (see Methods and Supplementary Table 1 for details). *, $P < 0.05$ vs AS, by one-way ANOVA plus Newman-Keuls post hoc test. n.s. = non-significant. Error bars are SEM.

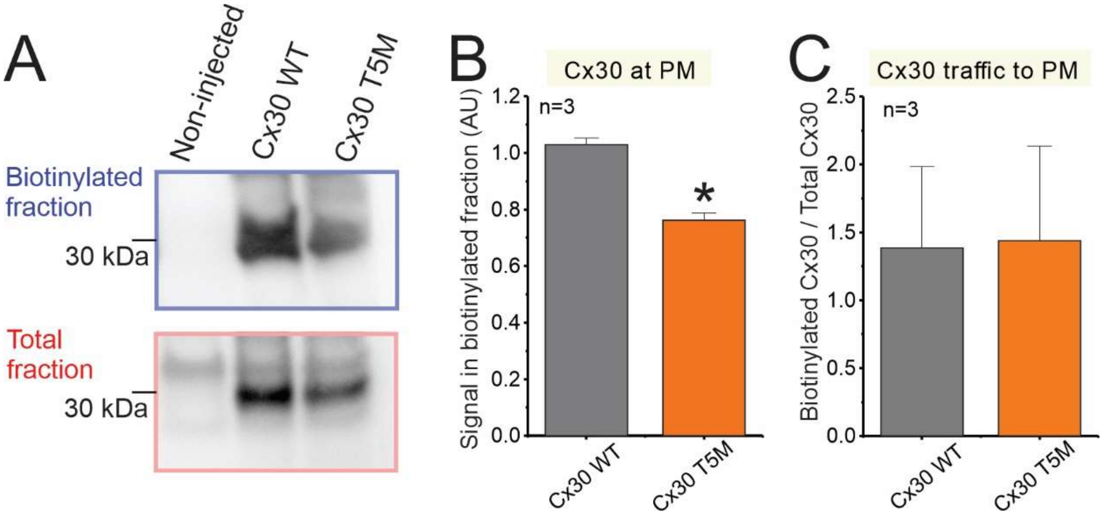


**Supplementary Fig. 12:** Human mutation T5M in Cx30 hemichannels reduces total protein levels but does not affect the traffic of hemichannels to the plasma membrane. **(A)** Representative Western blot of biotinylated fraction (i.e., the fraction containing plasma membrane proteins) and total fraction (i.e., the fraction obtained from the cell lysate before separation of biotinylated proteins). Oocytes were injected with WT or mutant Cx30 RNA. After 2 days, oocytes were treated with biotin to label and separate plasma membrane proteins. Antibodies targeting Cx30 were used to develop membranes (see Methods). Non-injected oocytes were used as a control. Note that both Cx30 WT and Cx30 T5M were found in the plasma membrane. **(B)** Quantification of the Western blot bands from the biotinylated fraction suggests that the T5M reduces the protein localization in the plasma membrane (PM). **(C)** Quantification of the ratio between biotinylated and total fraction indicates that T5M mutation reduces protein expression but do not affect Cx26 traffic to the plasma membrane (PM). Statistic was performed using one-way ANOVA plus Newman-Keuls post hoc test. Error bars are SEM.

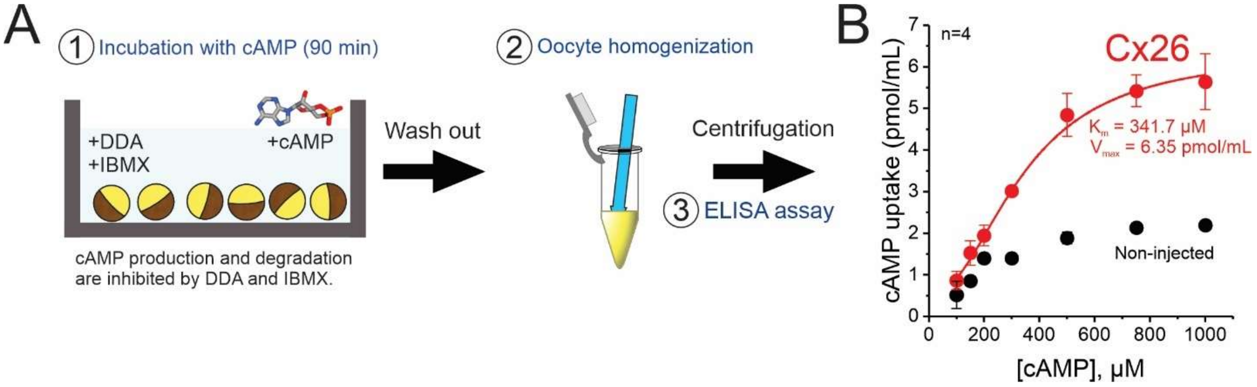


**Supplementary Fig. 13:** cAMP transport through Cx26 hemichannels is saturable. cAMP uptake was evaluated by an ELISA assay. **(A)** Scheme of the experimental approach. Oocytes expressing Cx26 WT were incubated for 90 min with a $Ca^{2+}$-free Ringer solution containing different cAMP concentrations. Endogenous production and degradation of cAMP were minimized by using DDA and IBMX (see Methods). After incubation, oocytes were homogenized, and cell lysates were centrifuged. Supernatant was used for cAMP measurement by ELISA following manufacturer instructions. Non-injected oocytes were used as control to determine endogenous Cx26-nonespecific cAMP uptake. **(B)** Quantification of cAMP uptake. Note that saturation is observed at the micromolar range. Error bars are SEM.

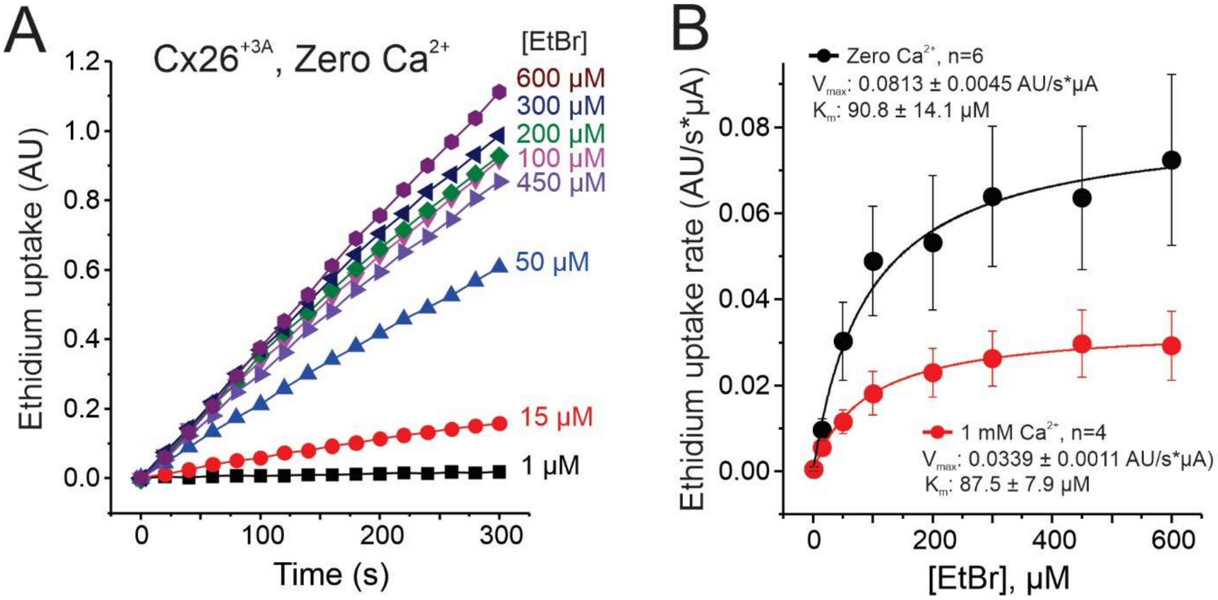


**Supplementary Fig. 14:** Low extracellular $Ca^{2+}$ increases ethidium transport through Cx26 +3A without affecting the apparent affinity. Ethidium permeability was evaluated using the TEVC/dye uptake assay (see Methods). **(A)** Representative time courses of ethidium uptake in oocytes incubated with different concentrations of ethidium bromide (EtBr). Measurements were performed in a $Ca^{2+}$-free Ringer solution. **(B)** Concentration-response curve for ethidium uptake in oocytes expressing Cx26 +3A. The ethidium uptake curve obtained in 1 mM extracellular $Ca^{2+}$ is shown to compare the effects of low extracellular $Ca^{2+}$, a gating modulator of Cx26 hemichannels, on permeation kinetics. Note that low $Ca^{2+}$ increases maximum transport but does not affect the Km. Data in **B** was normalized to the number of functional hemichannels using the magnitude of tail currents detected at -80 mV, after a voltage step pulse from -80 mV to 0 mV (40 s duration), and then fit with the Michaelis-Menten equation, to calculate Km and Vmax. Error bars are SEM.

| | Resting membrane potential (mV) | n | Conditions |
|---|---|---|---|
| **Non-injected** | -30.6 ± 2.6 | 20 | RS + 1 mM $Ca^{2+}$ |
| **Antisense (AS)** | -34.6 ± 1.9 | 29 | RS + 1 mM $Ca^{2+}$ |
| **Cx26 WT** | -14.0 ± 0.6 | 70 | RS + 1 mM $Ca^{2+}$ |
| **Cx26 +3A** | -14.9 ± 0.8 | 39 | RS + 1 mM $Ca^{2+}$ |
| | -5.3 ± 1.5 | 6 | RS, Zero $Ca^{2+}$ |
| **Cx26 W3A** | -23.9 ± 3.0 | 9 | RS + 1 mM $Ca^{2+}$ |
| **Cx26 D2N** | -21.5 ± 2.5 | 9 | RS + 1 mM $Ca^{2+}$ |
| **Cx26B** | -22.9 ± 3.7 | 9 | RS + 1 mM $Ca^{2+}$ |
| **Cx26B +2C** | -25.6 ± 2.0 | 16 | RS + 1 mM $Ca^{2+}$ |
| **Cx26 N14Y** | -16.3 ± 1.2 | 9 | RS + 1 mM $Ca^{2+}$ |
| **Cx26 S17F** | -22.4 ± 1.6 | 11 | RS + 1 mM $Ca^{2+}$ |
| **Cx30 WT** | -22.7 ± 2.6 | 13 | RS + 1 mM $Ca^{2+}$ |
| **Cx30 T5M** | -20.2 ± 0.9 | 10 | RS + 1 mM $Ca^{2+}$ |

**Supplementary Table 1:** Resting membrane potentials of oocytes used for dye uptake assays. All dye uptake recordings were performed using Ringer solution (RS) + 1 mM $CaCl_2$, with exception of data shown in Supplementary Fig. 14, which were partially performed under nominally zero $Ca^{2+}$ conditions. Values are Mean ± SE.
